# Regulatory landscapes and structural choreography of transcription initiation in spirochetes

**DOI:** 10.64898/2026.02.09.704077

**Authors:** Vilma K. Trapp, Tingting Wang, Tarek Hilal, Janne J. Makinen, Juho Kotikoski, Paavo J. Tavi, Ville Levola, Sari Paavilainen, Bernhard Loll, Markus C. Wahl, Georgiy A. Belogurov

## Abstract

Spirochaetota is a deeply rooted bacterial phylum of evolutionary and medical importance. We show that the spirochetal RNA polymerase (RNAP) has a diminished ability to melt promoters compared to its *Escherichia coli* counterpart. This deficiency is compensated by CarD and is not linked to the phylum’s natural resistance to the antibiotic rifampicin. CryoEM analysis revealed that the spirochetal RNAP disengages the −35 element during promoter unwinding, which contrasts with other bacterial RNAPs that release this element later during the initial transcription. Our research also details the unusually tight, non-sequence-specific binding of the spirochetal RNAP holoenzyme to DNA, which may be linked to the highly extended structure of the spirochetal nucleoid. We support these observations with a structural analysis of the dimeric holoenzyme non-specifically bound to DNA. Our results highlight the functional diversity of bacterial transcription systems and lay the foundation for understanding transcription regulation in pathogenic spirochetes.

## Introduction

RNA polymerase (RNAP) carries out an essential and most regulated step in gene expression, the synthesis of an RNA copy of the template DNA. All cellular RNAPs are multi-subunit enzymes that share homologous catalytic cores. Bacteria possess a single RNAP that is composed of five core subunits (ααββ’ω) ^1^. This multi-subunit core is guided to promoters by a dissociable subunit known as sigma (σ) factor ^2^. A σ factor binds core RNAP subunits forming a holoenzyme complex. A holoenzyme containing the primary σ factor initiates transcription at promoters that contain conserved sequence elements centered at −35 and −10 relative to the transcription start site (TSS). However, most bacteria also possess alternative σ factors directing the RNAP core to specific subsets of promoters.

In the widely studied *E. coli* system, the primary RNAP holoenzyme melts promoter DNA very efficiently ^3^. Consequently, regulation of promoter activity primarily occurs during recruitment ^4,5^ or through negative modulation of promoter melting by transcription regulator DksA ^6,7^ (**Fig. 1A**). In contrast, primary holoenzymes from Mycobacteria have reduced DNA melting capacities compared with the *E. coli* system ^8–10^ and necessitate allosteric activators CarD ^10–12^ and RbpA ^13–15^ for efficient melting of promoters. Holoenzymes containing alternative σ factors also rely on positive allosteric regulators to facilitate promoter DNA melting ^16,17^.

**Fig. 1.**
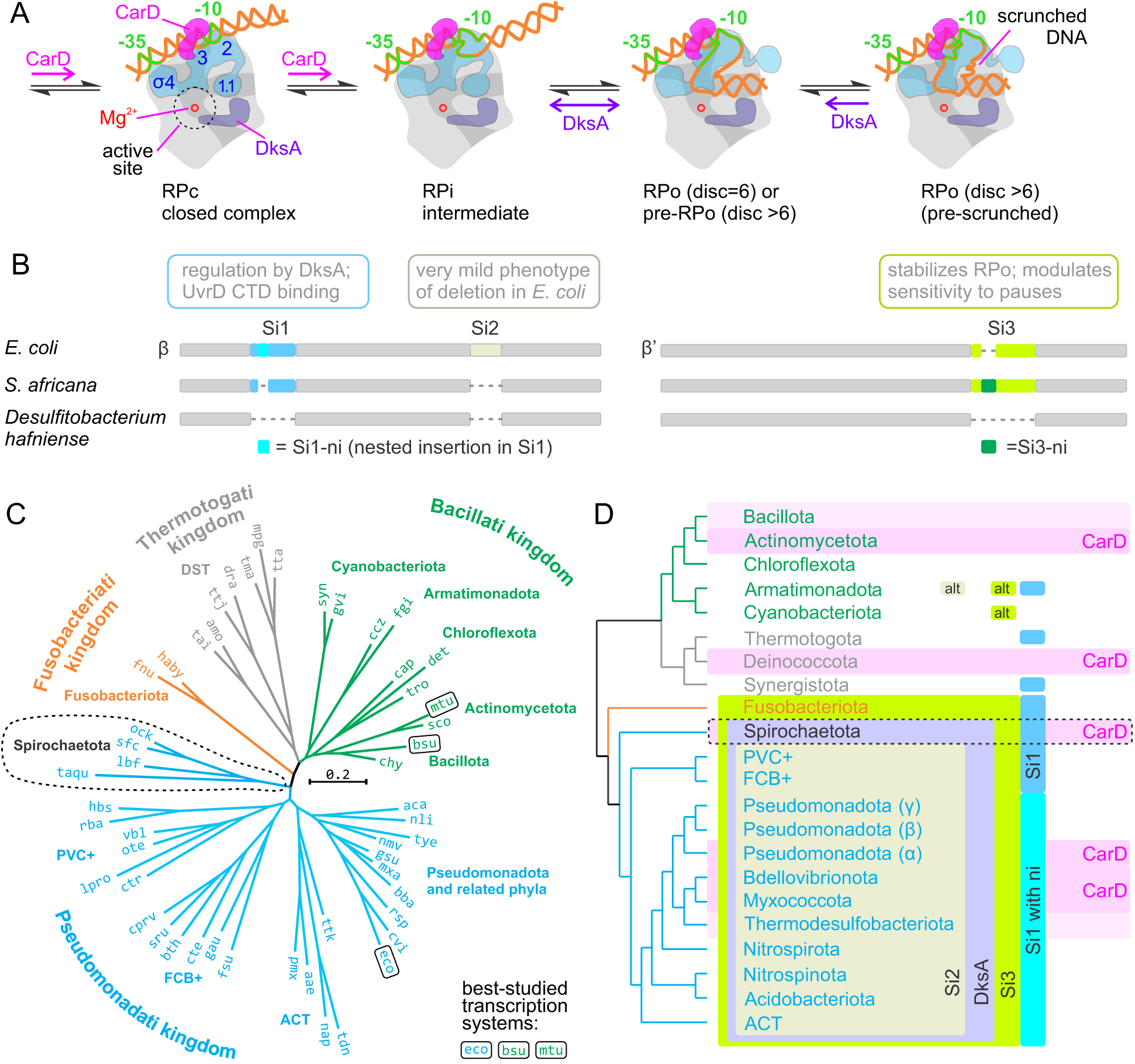
Spirochaetota is a deeply rooted phylum featuring a unique combination of lineage-specific inserts in RNAP and transcription initiation factors. (**A**) A simplified and abbreviated schematics of promoter recognition and melting by the primary RNAP holoenzyme. CarD helps anchoring the upstream DNA and facilitates melting of the −10 promoter element. DksA modulates the downstream DNA loading and promoter pre-scrunching. (**B**) The schematics depicting the location of Si1, Si2 and Si3 inserts in the primary sequences of β and β’ subunits of *E. coli* and *S. africana* RNAPs. *E. coli* Si1 and *S. africana* Si3 contain nested inserts, Si1-ni and Si3-ni, respectively. Insert-less RNAP from Bacillota is shown as a reference. (**C**) Bayesian molecular phylogenetic tree inferred using alignment of concatenated RNAP subunits and NusA transcription factor. Species names are described in Supplementary **Table 1**. (**D**) The distribution of lineage specific inserts and transcription initiation factors CarD and DksA. The presence of factors is designated only for phyla where CarD and DksA are highly prevalent. Light pink shading: CarD is frequently found but not prevalent. “alt” signifies the presence of sequence inserts that positionally correspond but are not homologous to Si2 and Si3 inserts of *E. coli* RNAP.

It is not precisely known why the primary holoenzyme from *E. coli* melts promoters more efficiently than its counterparts from Bacilli (e.g. *Bacillus subtilis*) and Mycobacteria (e.g. *Mycobacterium tuberculosis*). The difference is not attributable to the σ subunit ^10,18^ but may be linked to a lineage-specific insertion 3 (Si3) in the β’ subunit that enhances the stability of open promoter complexes (RPo) formed by *E. coli* holoenzyme^19^. Lineage-specific insertions are additional domains that are present in some RNAPs and not others (**Fig. 1B**). Both Si3 and the lineage-specific sequence insertion 1 (Si1) are also important for the modulation of the promoter melting by transcription regulator _DksA_ ^2^0–2^3^.

To broaden our understanding of transcription initiation in bacteria we performed a detailed biochemical and structural characterization of the RNAP holoenzyme from *Spirochaeta africana*, a free-living bacterium from the phylum Spirochaetota. This diderm bacterial phylum represents the most basal lineage within the Pseudomonadati kingdom (**Fig. 1C**) ^24^ and is notable for a unique combination of basal transcription factors and lineage-specific insertions in RNAP subunits (**Fig. 1D**). Spirochaetota are also characterized by their distinctively long and thin morphology and a periplasmic flagellum. A key unresolved question in spirochetal biology is how cellular components, including the transcription apparatus, are trafficked along their highly elongated body (∼1:100 width-to-length ratio).

We show by biochemical analysis that the spirochetal holoenzyme has diminished ability to melt promoters compared to *E. coli* and displays unusually tight non-sequence-specific association with DNA. We present cryoEM-based structural analyses of *S. africana* RNAP that reveal novel DNA binding modes: a disengagement of the −35 promoter element upon RPo formation and a non-sequence-specific engagement of the DNA minor groove by a dimeric holoenzyme. These findings broaden our understanding of bacterial transcription, offer insights into the mechanism of promoter escape and suggest phylum-level variations in how transcription machinery navigates the bacterial nucleoid.

## Results

### *In vitro* reconstituted transcription system of *S. africana*

Spirochetes processes both positive and negative regulators of ribosomal RNA transcription, CarD and DksA, respectively ^25^. Both CarD and DksA are measurably transcribed in *S. africana* (**Supplementary Fig. 1A**). While Spirochaetota are not unique in possessing both factors, the interplay of the two factors has not been investigated in detail previously. We cloned and heterologously expressed *S. africana* RNAP, primary σ factor, CarD and DksA in *E. coli*. We hand-picked six *S. africana* promoters of various strengths (**Supplementary Fig. 1B**) and constructed transcription templates (**Fig. 2A**).

**Fig. 2.**
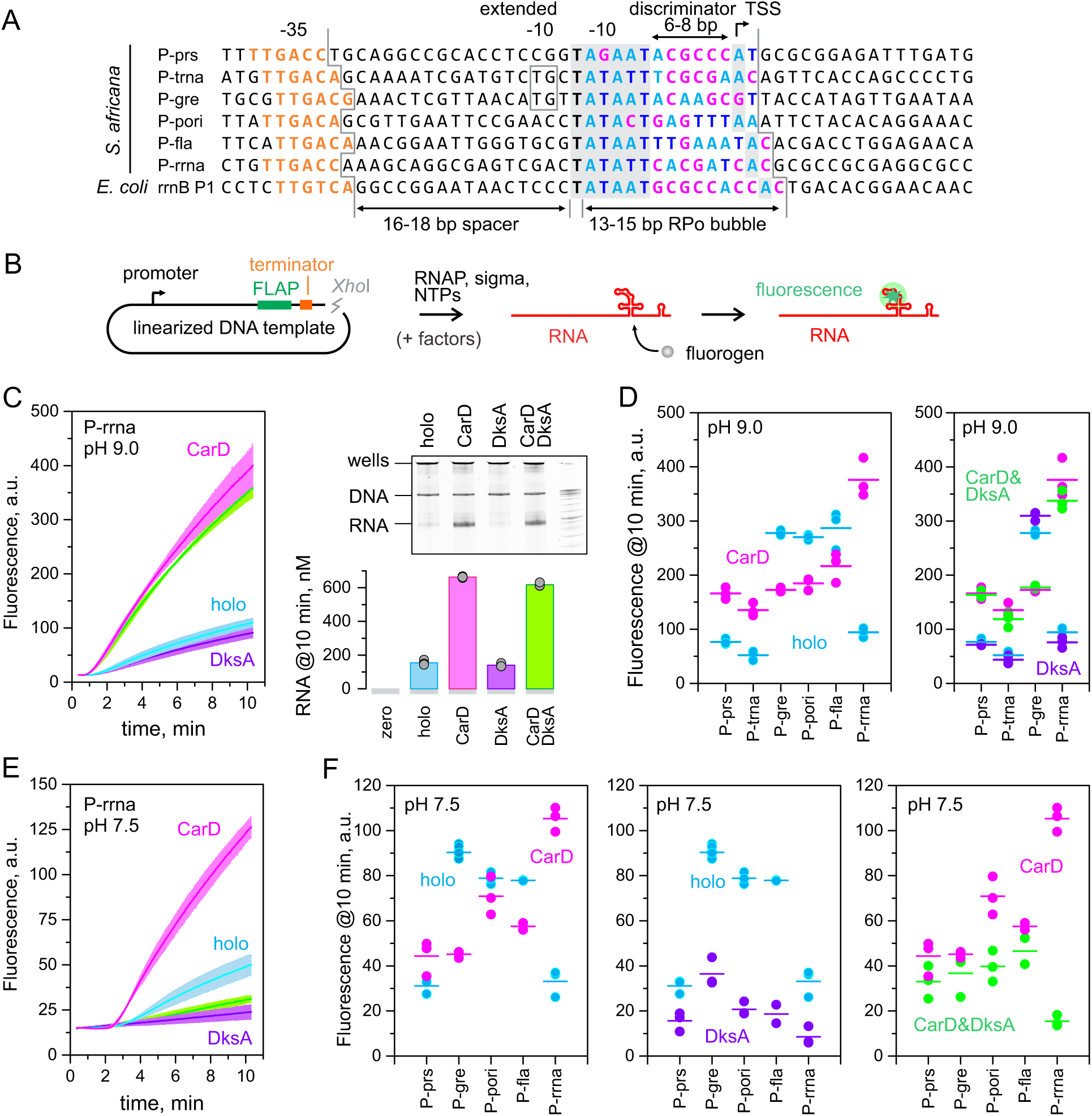
The effects of CarD and DksA on transcription initiation by *S. africana* RNAP holoenzyme. **(A)** Sequences of promoters used in this study aligned by −10 elements. **(B)** The schematic of the multi-round transcription assay using fluorescent light up aptamer (FLAP). **(C)** Transcription activity of the P-rrna promoter at pH 9.0. Real-time monitoring of transcription using FLAP (left graph), RNA concentration measured by Qubit assay (bar graph), SYBR-Gold stained denaturing PAGE gel (at 10 min). **(D)** Effects of CarD, DksA and their combination on transcription activity of six spirochetal promoters at pH 9.0. **(E)** Real-time monitoring of transcription from the P-rrna promoter at pH 7.5. **(F)** Effects of CarD, DksA and their combination on transcription activity of five spirochetal promoters at pH 7.5. Transcription assays were performed at 37 °C using 1 µM RNAP, 4 µM σ, 50 nM DNA template and where indicated 10 µM CarD and/or DksA. Bars in (D) and (F) indicate averages of three independent experiments.

We then used the above components to reconstitute multi-round transcription by *S. africana* RNAP *in vitro*. In a small subset of experiments for comparative purposes we also employed RNAP purified directly from *S. africana* cells by a combination of chromatographic techniques.

*S. africana* is a fermentative halo-alkaliphilic bacterium that grows optimally at pH between 9 and 10 and is dependent on sodium for growth ^26^. Most membrane pumps are annotated as sodium pumps suggesting sodium is used as a primary coupling ion, and cytoplasmic pH regulation is absent or inefficient ^25^. Our early experiments revealed that transcription activity of *S. africana* RNAP at pH 9.0 is approximately threefold higher than at pH 7.5. Accordingly, most *in vitro* transcription experiments with *S. africana* RNAP were performed at pH 9.0, 11 mM MgCl2 and total ionic strength of ∼180 mM. Unlike *Borrelia* RNAP ^27^, both heterologously expressed and fully native *S. africana* enzymes did not require Mn^2+^ for activity (**Supplementary Fig. 2**).

To monitor transcription activity, we used a multi-round assay where a preformed holoenzyme was added to a linearized plasmid encoding a transcription unit consisting of a spirochetal promoter followed by a 0.7 kb transcript encoding Broccoli fluorogenic light up aptamer (FLAP) upstream of the terminator (**Fig. 2B**) ^28^. In most experiments we used 50 nM of DNA template, 1 µM of holoenzyme and relatively strong promoters. In such systems, *S. africana* RNAP produces up to 600 nM RNA in 10 min allowing for monitoring of transcription output by measuring FLAP fluorescence, estimation of transcription yield using RNA-specific fluorescent dyes (Qubit fluorometer and kits), and visualization of the transcript in denaturing PAGE using SYBR Gold staining (**Fig. 2C**). FLAP fluorescence offered the best sensitivity and selectivity and was therefore employed as the main method throughout the study.

Throughout this study, we mainly report transcriptional activities in arbitrary fluorescence units, which suffice for the assessment of relative changes in transcription yield. Comparisons of FLAP and Qubit readings (**Fig. 2C**) suggest a conversion factor of 1.7 to express these units in nM of RNA. However, a portion of transcripts map outside the target transcription unit (**Supplementary Fig. 3**), implying a slightly smaller multiplier. Consistent with this, purified P-gre RNA, transcribed from a PCR-amplified short DNA template with minimal flanking DNA outside the transcription unit, indicates a coefficient of 1.4 for our instrumental setup ^28^. This coefficient allows for the estimation of absolute promoter activities in our *in vitro* assays. For instance, *S. africana* and *E. coli* holoenzymes initiate approximately once per minute at their cognate rRNA promoters at 37 °C. This is 60-fold lower than one initiation per second estimated for *E. coli* holoenzyme at rrnB P1 *in vivo* ^29^ possibly due to the presence of additional activators like Fis *in vivo* ^30^ as well as broader effects of genomic context.

### *S. africana* CarD stimulates promoters with suboptimal discriminators

We selected six promoters with diverse sequences and varying strengths by searching the *S. africana* genome with promoter patterns and considering *in vivo* transcript levels (**Supplementary Fig. 1B**). These promoters featured a TTGACN −35 element, a 16-18 bp spacer, a TANNNT −10 element, a 6-7 bp discriminator, and an A or G at TSS (**Fig. 2A**). We predicted TSS assuming initiation at a purine 6 bp from the −10 promoter element, or purine 7 bp away if the former was absent. We also mapped TSS by sequencing *in vitro* produced RNAs with Oxford Nanopore after ligating a 24 nt RNA adapter to the 5’ end. The RNA sequencing experiment confirmed the primary TSS but also indicated occasional slippage during transcription initiation at 5’-AAA (P-pori) and 5’-ACA (P-trna, P-fla) sequences (**Supplementary Fig. 4**).

*S. africana* RNAP initiated relatively efficiently at the P-gre, P-pori, and P-fla promoters irrespective of the presence of CarD (**Fig. 2D**). We designated these promoters as robust. CarD attenuated robust promoters up to 1.5 fold. *S. africana* holoenzyme alone initiated inefficiently at P-prs, P-trna, P-rrna but in the presence of CarD the activity of these three promoters was same or higher than that of robust promoters. We designated these promoters as tunable. While most measurements were performed at 10 µM CarD to ensure saturation, 1 µM of CarD was sufficient for the full effect of CarD at P-gre and P-rrna promoters (**Fig. 3A**).

**Fig. 3.**
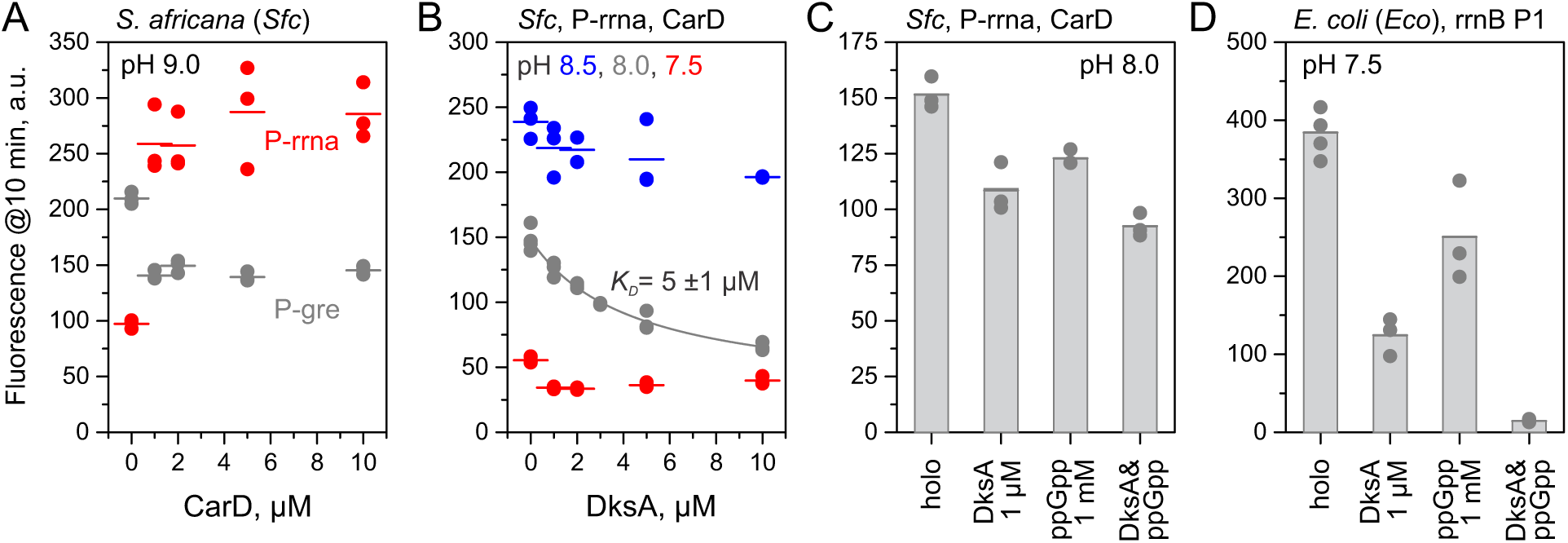
Concentration dependence of CarD and DksA effects on transcription activity of *S. africana* RNAP holoenzyme. **(A)** Modulation of P-gre and P-rrna activity by CarD as a function of CarD concentration. **(B)** The effect of DksA on transcription from the P-rrna promoter as a function of DksA concentration. CarD was present at 10 µM. The gray line is the bestfit to the hyperbolic equation. **(C)** Additive effect of ppGpp and DksA on transcription from the P-rrna promoter at pH 8.0. CarD was present at 2 µM. **(D)** Synergistic effect of ppGpp and *E. coli* DksA on transcription from rrnB P1 by *E. coli* holoenzyme at pH 7.5 (control experiment). Transcription assays were performed at 37 °C using 0.5 µM RNAP, 10 µM (in A-B) or 4 µM (in C-D) σ, 50 nM DNA template. Bars indicate averages of three independent experiments.

Robust promoters feature discriminators that are more purine and AT-rich than those of tunable promoters, suggesting that the former are easier to melt than the latter (**Fig. 2A**). At the same time, CarD effects did not correlate with the spacing between the −35 and - 10 elements, as it activated P-rrna with a consensual 17 bp spacer while attenuating P-pori that also features a 17 bp spacer. Similarly, the presence or absence of an extended - 10 element did not determine CarD action; P-trna (with the element) was activated, whereas P-gre (also with the element) was attenuated. Overall, the spirochetal holoenzyme closely resembled the properties of its mycobacterial counterpart ^10–12,31^. Adhering to its established role as an rRNA transcription regulator in Mycobacteria ^31^, spirochetal CarD significantly enhanced transcription from a spirochetal rRNA promoter.

### *S. africana* DksA is a pH-dependent negative regulator of transcription

When tested at pH 9.0, *S. africana* DksA did not affect transcription initiation at four tested promoters, regardless of the presence of CarD (**Fig. 2D**). However, DksA inhibited transcription 2-3-fold at all six promoters when tested at pH 7.5 (**Fig. 2E-F**). When present, CarD protected four out of six promoters from inhibition by DksA, but P-rrna and to some extent P-pori remained sensitive to DksA. As a result, in the presence of CarD, DksA specifically inhibited P-rrna and P-pori promoters. At pH 7.5, 1 µM of DksA added to 0.5 µM holoenzyme was sufficient for the maximal extent of inhibition of P-rrna transcription (**Fig. 3B**). A higher concentration of DksA (5-10 µM) was required to inhibit P-rrna twofold at pH 8.0, and at pH 8.5, the DksA effect on P-rrna transcription was mostly absent. Additionally, the alarmone ppGpp did not stimulate inhibition of P-rrna by DksA at pH 8.0 (**Fig. 3C**), although ppGpp did strongly stimulate inhibition of transcription from rrnB P1 by *E. coli* RNAP with *E. coli* DksA in a control experiment (**Fig. 3D**). Overall, our data indicate that *S. africana* DksA is a potent negative regulator of transcription initiation, that in the presence of CarD acts only at certain promoters, such as the rRNA promoter. Although *S. africana* does not grow below pH 8.0, a culture pre-grown at pH 9.2 remains viable for several days at pH 7.5 at 25 °C: most cells retain their spiral shape, stay motile and resume growth when transferred back to the alkaline medium (**Supplementary Movie 1**). These observations indicate that the transcription activity of *S. africana* DksA at pH 8 and below may be physiologically relevant.

### Stimulation by CarD is not linked to rifampicin resistance

Rifampicin resistance is a distinctive characteristic of spirochetal RNAP ^32^. This led us to investigate whether this resistance contributes to inefficient initiation at certain promoters, making spirochetal RNAPs reliant on CarD. Bioinformatic analysis suggested that rifampicin resistance in spirochetes is due to the strict conservation of the Asn residue (βAsn490 in *S. africana* RNAP) at a specific position in the β subunit ^33^. In contrast, most other bacterial phyla have a Ser residue at this position (βSer531 in *E. coli* RNAP). To test this idea, we constructed a βN490S variant of *S. africana* RNAP and a βS531N variant of *E. coli* RNAP and examined their sensitivity to rifampicin. Wild-type *S. africana* RNAP was completely resistant to 1 µM rifampicin, whereas transcription by the βN490S variant was strongly inhibited by 1 µM rifampicin (**Fig. 4A**). Conversely, 1 µM rifampicin completely abolished transcription by the wild-type *E. coli* RNAP but had no effect on transcription by the βS531N variant (**Fig. 4B**). These results demonstrate that the phylum-specific Asn residue is the primary determinant of rifampicin resistance in Spirochaetota.

**Fig. 4.**
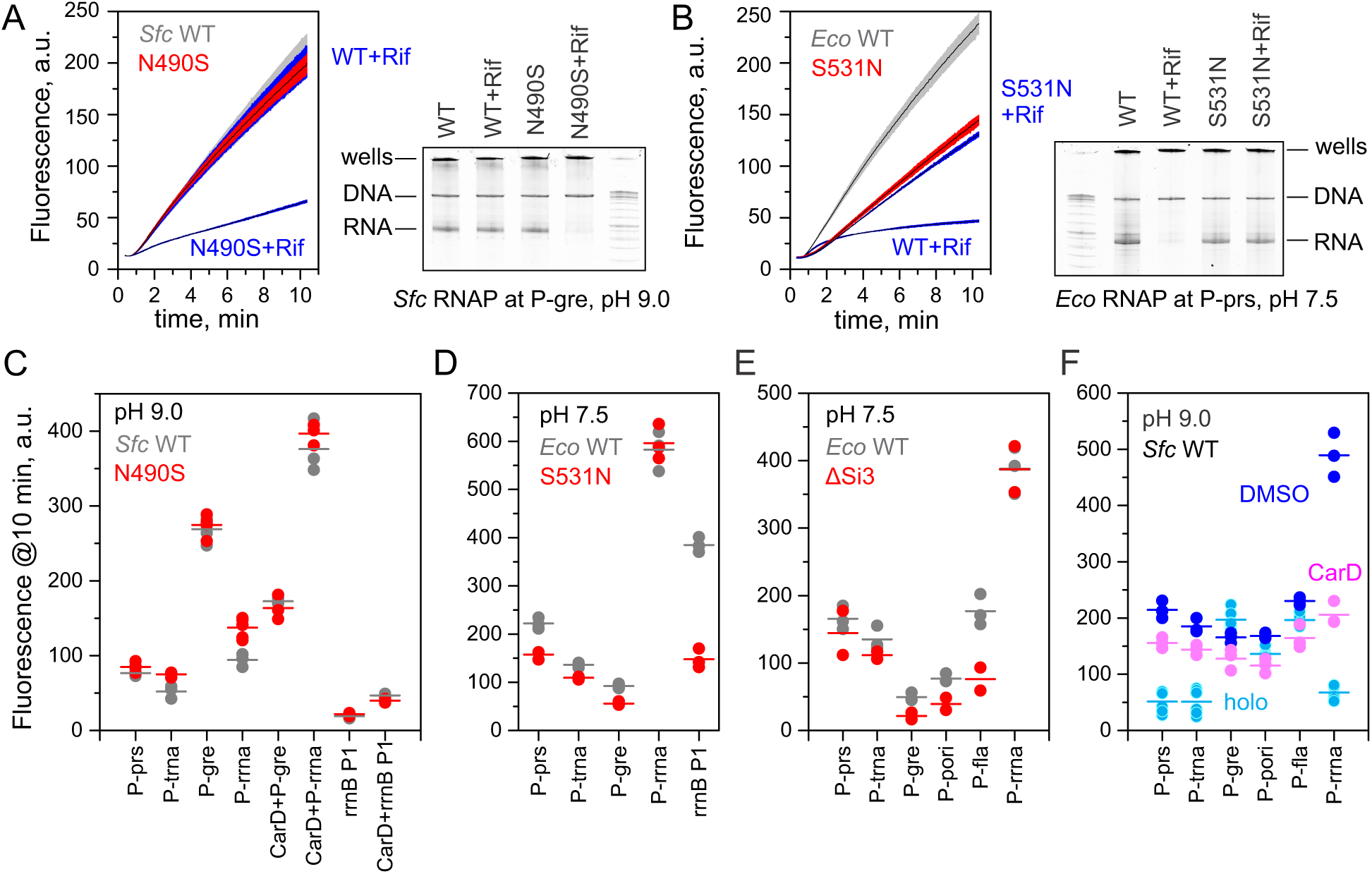
Resistance to rifampicin does not govern the promoter preferences of *S. africana* RNAP. **(A)** The effect of 1 µM of rifampicin on transcription by the wild-type and βN490S variant of *S. africana* holoenzyme at P-gre promoter. Left: real-time monitoring of transcription using FLAP assay. Right: transcription reactions were incubated for 10 min, reaction products were resolved on 6% denaturing PAGE gel and stained with SYBR-Gold. **(B)** The effect of 1 µM of rifampicin on transcription by the wild-type and βS351N variant of *E. coli* holoenzyme at P-prs promoter. **(C)** The effect of βN490S substitution on the promoter preferences of *S. africana* holoenzyme. **(D)** The effect of βS531N substitution on the promoter preferences of *E. coli* holoenzyme. **(E)** The effect of deletion of Si3 on the promoter preferences of *E. coli* holoenzyme. **(F)** The effect of 10% DMSO on the promoter preferences of *S. africana* holoenzyme. Transcription assays were performed at 37 °C using 1 µM RNAP, 4 µM σ, 50 nM DNA template at pH 9.0 (*S. africana*) and pH 7.5 (*E. coli*). CarD was added at 10 µM where indicated. Measurements in (E) and (F) were performed using end-point-only FLAP assay. Fluorescence readings were scaled × 1.5 fold to account for the dilution during quenching by fluorogen-EDTA mixture. Bars indicate averages of three independent experiments.

We then compared the performance of the wild-type and variant holoenzymes on a set of four *S. africana* promoters. The results showed a negligible effect of βN490S on *S. africana* holoenzyme activity, with only marginal changes observed at P-rrna and P-trna promoters (<1.5 fold) (**Fig. 4C**). Additionally, the βN490S substitution did not impact the effects of CarD at P-rrna and P-gre promoters. Similarly, the βS531N substitution had a minimal effect on *E. coli* holoenzyme activity at *S. africana* promoters (**Fig. 4D**). However, βS531N substitution significantly (3-fold) impaired the ability of *E. coli* holoenzyme to initiate at *E. coli* rrnB P1.

### CarD aligns promoter preferences of *S. africana* and *E. coli* holoenzymes

Next, we compared the activity of *S. africana* and *E. coli* holoenzymes using a full set of six *S. africana* promoters. To speed up data acquisition, we used an end-point-only implementation of the FLAP assay. Transcription reactions were performed for 10 min in a thermoblock, quenched with a fluorogen-EDTA mixture followed by fluorescence measurements. In the absence of CarD, the relative promoter preferences of *E. coli* (**Fig. 4E**, gray) and *S. africana* holoenzymes (**Fig. 4F**, cyan) were significantly different. *E. coli* holoenzyme displayed high activity at tunable promoters (P-prs, P-trna, P-rrna) where the basal activity of *S. africana* holoenzyme is low but performed poorly at robust promoters (P-gre, P-pori) where the basal activity of *S. africana* holoenzyme is high.

Only P-fla displayed high activity with both holoenzymes. CarD made *S. africana* holoenzyme more similar to its *E. coli* counterpart by activating tunable promoters and slightly attenuating robust promoters (**Fig. 4F**, magenta). Interestingly, 10% DMSO, known to facilitate DNA melting, closely mimicked the effect of CarD on the spirochetal holoenzyme across all promoters and established even better alignment of promoter preferences between *S. africana* and *E. coli* holoenzymes (**Fig. 4F**, blue). These observations suggest that the spirochetal holoenzyme has intrinsically lower capacity to melt promoters than the *E. coli* enzyme, and this deficiency is compensated by CarD.

We then investigated whether a modified *E. coli* holoenzyme lacking Si3 inserts, known to form an unstable RPo ^19^, would exhibit reduced activity at tunable promoters. Contrary to the expectations, ΔSi3 *E. coli* holoenzyme displayed similar promoter preferences and absolute activity to the wild-type *E. coli* holoenzyme (**Fig. 4E**). These findings indicate that the Si3 insert and RPo stability do not significantly influence transcription initiation in our high-NTP assays (1 mM of each NTP). Therefore, differences in promoter melting capacity between *S. africana* and *E. coli* RNAP are related to the forward rate of promoter melting ^11^, not the stabilization of the melted state.

### Structure of *S. africana* RPo: the transcription bubble and CarD

To illuminate the structural choreography of transcription initiation in spirochetes we determined the cryoEM structures of promoter complexes formed by *S. africana* RNAP holoenzyme at P-rrna and P-gre promoters in the presence of CarD at the overall resolution of about 3 Å (**Supplementary Table 1**). For the P-gre promoter, we also determined the structure in the absence of the factor. Unsurprisingly, the overall structures of *S. africana* RPo closely resembled the structures of other bacterial RPo. The single-stranded ntDNA could be traced in all three structures. In contrast, the density for the single-stranded tDNA in the RNAP cleft was mostly absent in P-rrna and P-gre RPo without CarD but could be fully traced in P-gre RPo with CarD (**Supplementary Fig. 5**). CarD binds to RNAP in canonical conformation closely resembling the binding mode previously observed in Mycobacteria and Deinococcota ^34–36^ (**Fig. 5A**). Specifically, CarD binds to a protrusion with its N-terminal tudor domain and inserts a conserved Trp of its C-terminal α-helical domain into the minor groove just upstream of the −10 promoter element. The same binding pose was observed at both CarD-stimulated P-rrna and CarD-attenuated P-gre promoters. Overall, spirochetal CarD is well positioned to promote melting of the −10 element, secure the upstream DNA binding to the holoenzyme and indirectly facilitate further steps in promoter melting.

**Fig. 5.**
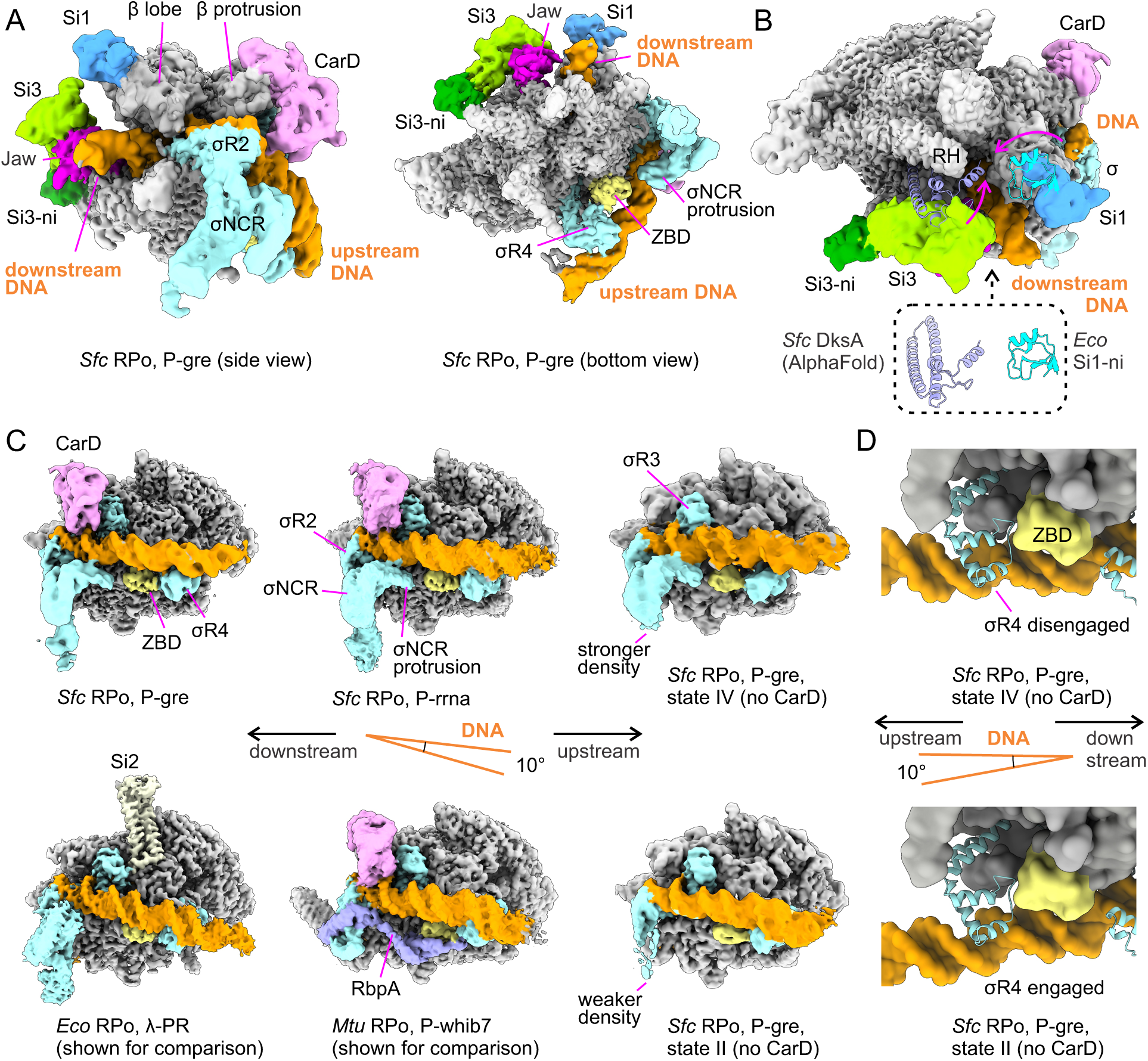
Structures of the *S. africana* RPo at P-gre and P-rrna promoters. **(A)** CarD binds to β protrusion and upstream DNA in a canonical conformation previously observed for mycobacterial CarD. Si3 insert positions the Jaw domain to interact with the downstream DNA. **(B)** Si1 is positioned far away from all other RNAP domains. The *E. coli* Si1-ni subdomain (blue cartoon) and *S. africana* DksA (purple cartoons) are modeled-in based on the alignment with *E. coli* holoenzyme structures. Magenta arrows indicate motions of lobe and Si3 domains that bring *Eco* Si1-ni in contact with either Si3 (PDB ID 5IPN) or DksA (PDB ID 7KHI). RH, β’ Rim Helices. **(C)** The overall course of the upstream DNA in *S. africana* RPo. *E. coli* (PDB ID 7MKD) and *M. tuberculosis* (PDB ID 7KIN) RPo are shown for comparison. The top row displays structures where the −35 promoter promoter element is disengaged from σR4, whereas the bottom row displays canonical engaged conformations. **(D)** Close-up view of the disengaged (top) and engaged (bottom) conformations of the upstream DNA in *S. africana* RPo. Structures in (C) and (D) were aligned using the β subunit as a reference. The central schematics highlight the differing paths of the upstream DNA between the top and bottom structures.

### Structure of *S. africana* RPo: lineage-specific inserts

CryoEM reconstructions of *S. africana* RPo revealed the structural fold of major lineage-specific inserts Si1 and Si3 (**Fig. 5A**). The structures showed that spirochetal Si1 is a substructure of *E. coli* Si1 and features the same overall fold. Spirochetal Si1 is missing the subdomain (named Si1-1.2 by Parshin *et al.*) that has been implicated in functioning of *E. coli* DksA ^21^. For consistency, we refer to Si1-1.2 as Si1-ni, for nested insertion. *E. coli* DksA influences promoter melting by inserting itself between Si1, Si3 and the Rim Helices (RH), thereby changing the mobility of the β lobe domain that in turn affects the σR1.1 ejection and promoter melting ^23^. *E. coli* Si1-ni forms the bulk of the contact interface between DksA and Si1 (**Fig. 5B** and PDB ID 7KHI). The absence of Si1-ni in spirochetes may therefore prevent the Si1 from interacting with DksA, effectively decoupling the entire lobe from DksA. Nevertheless, our biochemical data show that *S. africana* DksA strongly affects transcription initiation *in vitro*. Furthermore, two large bacterial superphyla, PVC+ (Planctomycetota, Verrucomicrobiota, Chlamydiota and related phyla) and FCB+ (Fibrobacterota, Chlorobiota, Bacteroidota and related phyla), feature DksA and Si1 without Si1-ni (**Fig. 1C-D**). These observations suggest that while *E. coli* DksA evolved to depend on the Si1-ni for functioning, DksA from Spirochaetota, PVC+ and FCB+ utilize rudimentary mechanisms that do not require Si1-ni.

*S. africana* Si3 has the same overall fold as *E. coli* Si3, with the *E. coli* insert being a substructure of the *S. africana* insert. *S. africana* Si3 incorporates an additional phylum-specific domain that we named Si3-ni, for nested insertion (**Fig. 5A-B**, dark green). Si3-ni rests on the β’ Shelf region and is distant from the RH, Jaw, and Si1. Therefore, the presence of Si3-ni does not compensate for the absence of Si1-ni. *E. coli* Si3 positions the Jaw domain ^37,38^ to interact with downstream DNA in RPo. Interestingly, although the Jaw domain is present in all bacterial RNAPs, it only interacts with downstream DNA when positioned by the canonical Si3 (e.g. PDB ID 7MKD). In RPo formed by Si3-less RNAPs (e.g. PDB IDs 7KIN, 7CKQ) or those with non-canonical Si3 (e.g. PDB ID 8GZG), the Jaw is completely disengaged from the downstream DNA. Our cryoEM reconstructions revealed that spirochetal Si3 positions the Jaw to interact with downstream DNA in a manner similar to that in *E. coli* RPo thereby reaffirming the trend described above (**Fig. 5A**).

### Structure of *S. africana* RPo: upstream DNA and σR4

A noticeable unique characteristic of *S. africana* RPo is the α-helical protrusion of the σNCR (σ non-conserved region) that extends towards the β subunit N-terminal zinc binding domain (ZBD) and contacts the phosphate backbone of the template DNA strand at registers −17 to −19 (**Fig. 5A-B**). As a result, the σR2-proximal upstream DNA duplex is almost completely encircled by the proteinaceous environment: tDNA −13 to - 15 is contacted by CarD, −17 to −19 by the σNCR protrusion, whereas ntDNA −14 to −17 is contacted by σR2 and σR3. Interestingly, the α-helical protrusion of the σNCR occupies a similar volume to that of the C-terminal region of the transcription factor RbpA in mycobacterial RPo (**Fig. 5C**). Thus, some of the RPo stabilizing functionality of mycobacterial RbpA (not present in spirochetes) could be encoded in the spirochetal σ itself in the form of α-helical protrusion of the σNCR.

The main chain atoms of σR4 domains align well in *S. africana* and *E. coli* RPo structures, and the amino acid sidechains implicated in the recognition of the −35 promoter element ^39^ are conserved (Arg561/584 Glu562/585, Gln566/589; *S. africana* and *E. coli* numbers respectively). However, closer examination of CarD-bound structures revealed that the recognition α-helix of σR4 interacts with the upstream DNA in an atypical and non-sequence-specific manner. σR4 usually interacts with the upstream DNA as a standard helix-turn-helix motif: the N-terminal tip of the recognition α-helix inserts halfway into the major groove and forms base-specific contacts with the −35 promoter element, while the positioning α-helix interacts with the phosphate backbone (**Fig. 5C-D**, bottom panels). In *S. africana* RPo, the spacer DNA slightly curves towards the β Flap and β’ Dock (**Fig. 5C-D**, top panels). This curving causes a significant shift of the upstream DNA duplex from its usual location when it approaches σR4: the entire recognition α-helix of σR4 is inserted into the major groove, potentially interacting with the phosphate backbone but unlikely forming base-specific contacts with the −35 promoter element. Consistently, both the upstream DNA and σR4 are positioned nearly identically in the structures of P-gre and P-rrna RPo featuring 16 bp and 17 bp spacers, respectively, suggesting the absence of sequence-specific contacts between σR4 and the −35 element. In the CarD-less RPo, the upstream DNA exhibits significant flexibility, allowing for reconstruction of four distinct structural states. Two of these states (II and III) feature the −35 element canonically engaged with σR4, while the other two (I and IV) display disengaged conformations, similar to those observed in CarD-bound RPo at P-gre and P-rrna (**Fig. 5C-D**, **Supplementary Movie 2**). Interestingly, the helical protrusion of σNCR appears partially disordered when the −35 element is canonically engaged (states II and III) (**Fig. 5C**, bottom right) but fully ordered when the upstream DNA is disengaged (all other structures of *S. africana* RPo). This suggests a connection between the upstream DNA disengagement and the helical protrusion of σNCR.

### Structure of *S. africana* holoenzyme dimer bound to DNA

During cryoEM structural analysis of *S. africana* holoenzyme at the P-gre promoter, using high concentrations of holoenzyme and DNA, we unexpectedly observed a complex featuring two holoenzyme protomers interacting with the central DNA duplex in a non-sequence-specific manner (**Fig. 6**). Each holoenzyme protomer additionally contains a DNA duplex end-bound in the downstream DNA channel: an *in vitro* artifact due to abundant DNA ends previously observed also in the *E. coli* system (PDB ID 7KHC). **Fig. 6A** illustrates the overall mode of holoenzyme engagement with the central DNA duplex, whereas schematic in **Fig. 6B** reveals the core architecture of the complex: two holoenzyme protomers dimerize via ZBDs and interact with the central duplex DNA via αCTD and the σR4 recognition helix. The two protomers are related through a 180-degree rotation around an axis that passes through the interface between the ZBDs and is perpendicular to the plane of the page. With the core architecture in mind, it becomes evident from the leftmost image in **Fig. 6B** that the space between the protomers is occupied by two interlocked σNCR domains, one from each protomer.

**Fig. 6.**
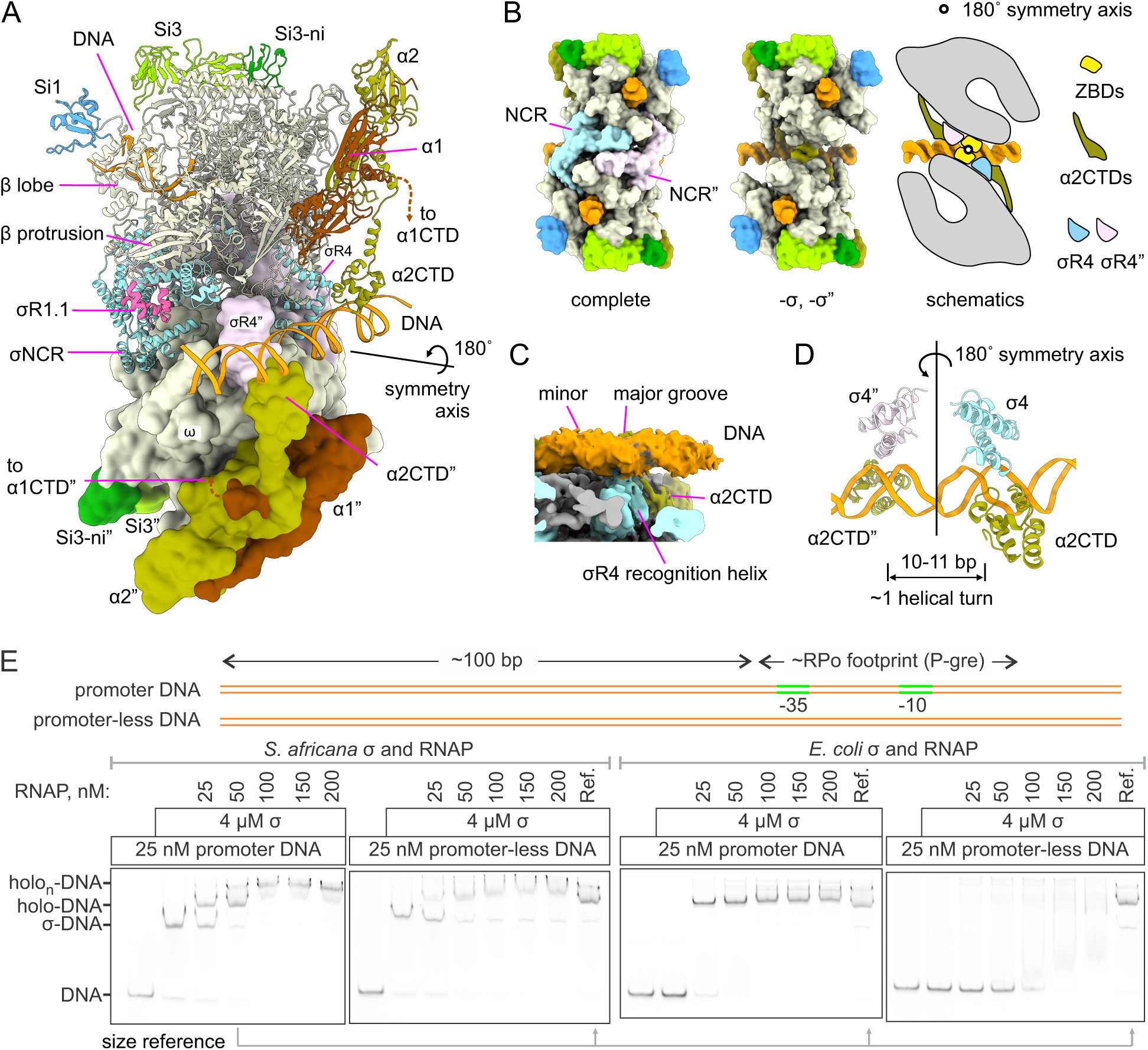
***S. africana* holoenzyme dimer bound to DNA in a non-sequence-specific manner. (A)** The top protomer and DNA duplexes are shown as cartoons, the bottom protomer is depicted as a surface. The σ subunits are colored light cyan (top protomer), and light pink (bottom protomer). **(B)** A view along the axis of the DNA end-bound within the downstream DNA channel. Both protomers are shown as surfaces. The left projection shows the complete structure. The middle projection omits σ subunits to reveal the juxtaposed ZBDs at the interface between the protomers. The right schematic visualizes the core architecture of the complex. **(C)** Cut-out view illustrating the engagement of the minor groove of DNA by σR4. **(D)** Close-up view illustrating the engagement of the minor DNA groove by σR4 and α2CTD. **(E)** Electrophoretic mobility shift assay (EMSA) was used to assess the binding of *S. africana* and *E. coli* holoenzymes to DNA fragments with and without a promoter at submicromolar concentrations of RNAP and DNA.

The most interesting feature of the complex is the mode of interaction with the central DNA duplex. The σR4 recognition helix of each protomer clearly engages the minor groove of the straight DNA duplex: an inherently non-sequence-specific type of interaction (**Fig. 6C**). Each protomer interacts with the central DNA duplex via a forked arm composed of σR4 and the α2CTD. The forked arm resembles the αCTD dimer interacting with the UP-element at *E. coli* rrnB P1 (PDB ID 7KHC), but the σ recognition helix is involved instead of the α1CTD, and the interaction with DNA takes place much closer to the RNAP body. The forked arms contributed by each protomer interact with the minor groove one helical turn apart (10-11 bp, **Fig. 6C**) and are related by a 180 degrees rotational symmetry as do the entire protomers. The same interaction can be principally formed by a monomeric holoenzyme using a single forked arm, albeit with a reduced affinity due to the loss of cooperativity between protomers.

Another interesting feature of the structure is an additional α-helical density observed in the volume bordered by σR2, the σNCR helical protrusion and σR3. In RPo this volume is occupied by the duplex DNA immediately upstream of the −10 element. We attributed this additional density to the σR1.1 globular domain (**Fig. 6A**, magenta cartoon). To verify this assignment, we probed the location of σR1.1 in *S. africana* holoenzyme by *in vitro* crosslinking with disuccinimidyl sulfoxide (DSSO) followed by mass spectrometry analysis (CLMS). We identified two distinct clusters of Lys residues that crosslink to Lys residues within σR1.1 (**Supplementary Fig. 6**). σR1.1-crosslinkable Lys residues in Switch 2 region, the Rudder Loop, and the Jaw domain suggest that σR1.1 is located in the downstream DNA channel, as observed in structures of *E. coli* holoenzyme ^40,41^ and early intermediates in the promoter melting pathway ^23,42^. At the same time, σR1.1-crosslinkable Lys residues in σR2, σR3, σNCR, and CarD suggest that σR1.1 is located in place of the upstream fork junction supporting our assignment of the additional density to σR1.1. Our CLMS analysis suggests that σR1.1 naturally alternates between locations in the downstream DNA channel and near σR3.

### *S. africana* holoenzyme efficiently binds DNA in a non-sequence-specific manner

Given that the dimeric holoenzyme complex with DNA used in structural analysis was assembled at a high concentration of RNAP and DNA, we used an electrophoretic mobility shift assay (EMSA) to investigate the binding of *S. africana* and *E. coli* holoenzymes to DNA at physiologically relevant concentrations. The promoter DNA fragment contained the P-gre promoter, 28 bp of DNA downstream of the TSS, and 101 bp of DNA upstream of the −35 element (**Fig. 6E**). Addition of 4 µM of *S. africana* σ to 25 nM of the promoter DNA fragment resulted in a distinct, slower-migrating band, that we interpreted as a σ-DNA complex. Subsequent addition of 25 nM RNAP led to an even slower-migrating band, that we interpreted as a promoter-bound holoenzyme-DNA complex (holo-DNA). This interpretation was supported by the formation of a similarly sized band when the DNA fragment was titrated with *E. coli* holoenzyme.

Increasing *S. africana* RNAP concentration to 100 nM caused the disappearance of σ-DNA and holo-DNA bands and the emergence of an even slower-migrating band, which we termed the multi-holoenzyme-DNA complex (holo_n_-DNA). This complex likely consists of a promoter-bound holoenzyme and additional holoenzyme(s) bound non-specifically upstream. In contrast, we did not observe formation of multi-holoenzyme-DNA complexes when titrating with *E. coli* holoenzyme. We also titrated a promoter-less DNA fragment with *S. africana* and *E. coli* holoenzymes and observed formation of distinct complexes in the former but not the latter case (**Fig. 6E**). These findings indicate that the *S. africana* but not *E. coli* holoenzyme possesses a strong tendency for non-sequence-specific association with DNA at physiological concentrations.

## Discussion

We report a comprehensive structural-functional analysis of transcription initiation in Spirochaetota. This diderm bacterial phylum is the most basal lineage within the Pseudomonadati kingdom and is markedly divergent from characterized model systems. We show that spirochetal transcription initiation displays a combination of features from both *E. coli* and Mycobacteria. Structurally, the spirochetal RPo is similar to that of *E. coli* and less so to Mycobacteria and Bacilli, consistent with sequence and phylogenetic analyses. In particular, spirochetes feature lineage-specific sequence inserts Si1 and Si3 that have been implicated in stabilizing RPo and regulation of transcription initiation by DksA in *E. coli*. However, biochemically, the spirochetal holoenzyme behaves more like the mycobacterial enzyme, exhibiting a weaker intrinsic ability to melt promoter DNA. This deficiency is overcome by the CarD activator, which is structurally similar to its mycobacterial counterpart and functions analogously in spirochetes. This differs from the situation in α-proteobacteria, where CarD has a more specialized function and primarily facilitates melting of promoters with non-canonical - 10 elements ^43^.

In contrast to mycobacteria, spirochetes possess a DksA regulator, similar to that found in *E. coli*. We demonstrate that *S. africana* DksA functions as a pH-responsive negative regulator of ribosomal RNA transcription but suggest that its overall regulatory repertoire might be reduced due to spirochetes lacking the Si1-ni subdomain that is important for the full functionality of *E. coli* DksA ^21^. In particular, inhibition of *S. africana* holoenzyme by DksA was not enhanced by ppGpp, which contrasts with the strong DksA-ppGpp synergy observed in the *E. coli* system. These results align with *in vivo* studies in *Borrelia*. Although ΔDksA *Borrelia* strains show deficiencies in adapting to nutrient-limited conditions, reduced virulence, and altered transcriptomes ^44–46^, DksA and ppGpp regulation appear to function independently, in parallel.

Resistance to rifampicin is a unique and conserved feature of spirochetal RNAPs, the functional significance of which remains unclear given that these species inhabit various environments and are not universally exposed to rifampicin-like compounds. In other bacterial phyla, acquiring rifampicin resistance often incurs a fitness cost ^47,48^. However, the rifampicin-sensitive βN490S RNAP displayed the same activity at spirochetal promoters as the wild-type *S. africana* RNAP. A converse βS351N change in the *E. coli* RNAP also did not affect its performance at spirochetal promoters. Interestingly, βS351N significantly reduced activity of *E. coli* RNAP at *E. coli*’s own rrnB P1, whereas *S. africana* holoenzymes exhibited very low activity at rrnB P1, both with and without CarD (**Fig. 4C-D**). These observations are consistent with the idea that spirochetes may utilize a limited set of promoters where rifampicin resistance does not impose a fitness cost, in contrast to *E. coli*’s broader range of promoters.

Our structural analysis revealed that the spirochetal holoenzyme disengages the −35 promoter element from σR4 upon promoter melting. This contrasts with other bacterial holoenzymes studied to date, which maintain interactions with both the −35 and −10 promoter elements during RPo formation. Recent structural studies of deinococcotal and mycobacterial RNAPs showed that −35 disengagement occurs late during the initial transcription via the allosteric displacement of the σ-finger (a loop region linking σR3 and σR4) by the extending RNA-DNA hybrid (**Fig. 7A**) ^49,50^. Notably, −35 disengagement precedes, and may be essential for the subsequent release of the −10 element in the mycobacterial system. In the spirochetal system, the coupling of −35 disengagement with promoter melting may facilitate the earlier release of the −10 promoter element during the initial transcription, leading to a promoter escape pathway with less DNA scrunching compared to other bacterial systems (**Fig. 7A**). This may contribute to the efficient initiation at consensus-like promoters such as P-gre and P-pori, where *S. africana* RNAP outperforms the *E. coli* counterpart.

**Fig. 7.**
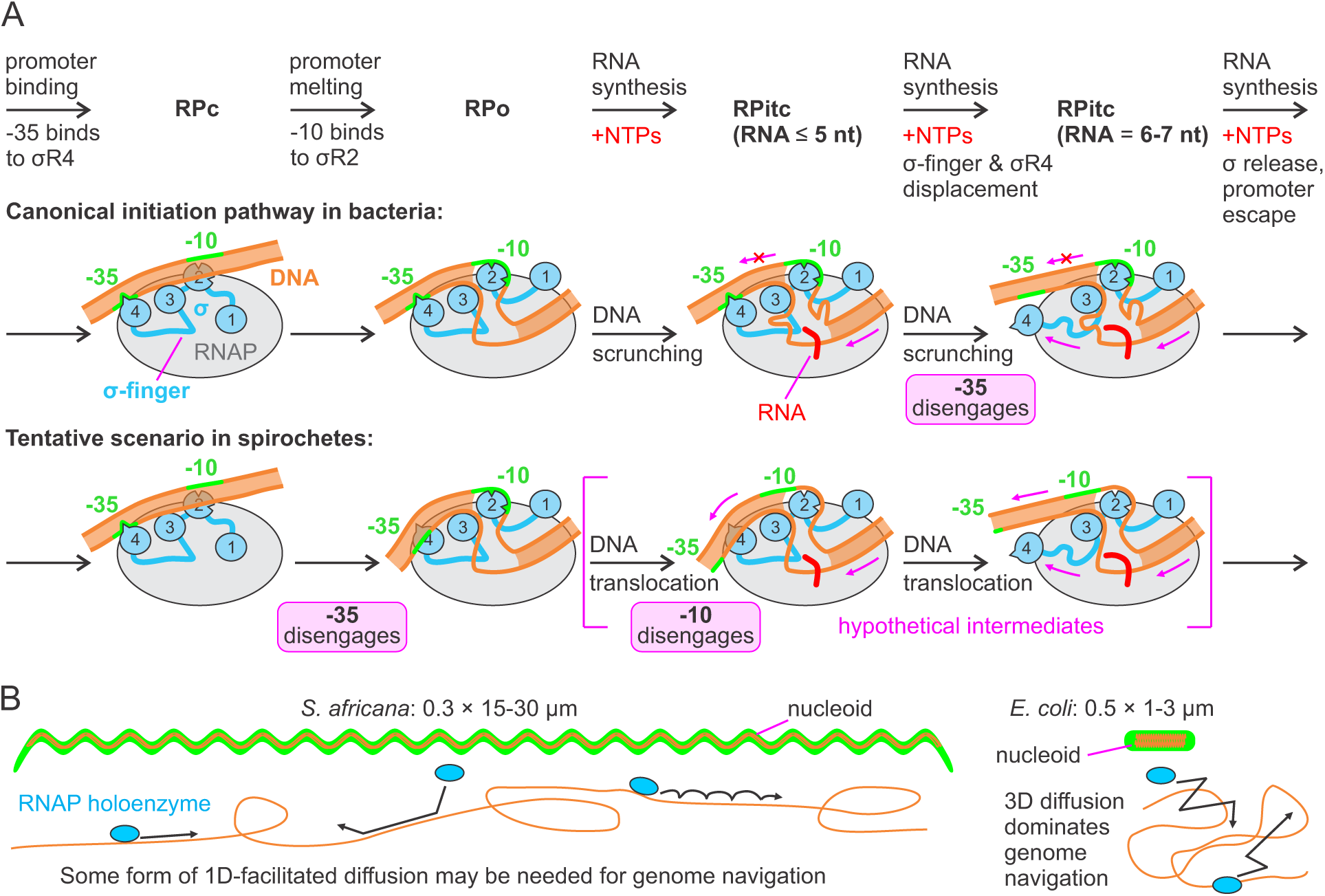
Distinctive features of the spirochetal transcription apparatus. **(A)** Bacterial RNAP holoenzymes typically disengage the −35 promoter element after synthesizing RNA six or more nucleotides long (top schematic). In the spirochetal system, the coupling of −35 disengagement with promoter melting may facilitate the earlier release of the −10 promoter element during the initial transcription, leading to a promoter escape pathway with less DNA scrunching (bottom schematic). Magenta arrows indicate the movement of DNA, σ-finger and σR4, or lack of movement (if crossed). **(B)** Schematic comparison of the cell morphology of *S. africana* and *E. coli* provides a rationale for the tighter non-sequence-specific association of the spirochetal holoenzyme with DNA.

The spirochetal holoenzyme displays an unusually strong, non-sequence-specific association with DNA and can bind to the minor groove (**Fig. 6**). Collectively, these findings suggest that the spirochetal holoenzyme predominantly exists in a DNA-associated form and navigates the genome via one-dimensional facilitated diffusion mechanisms, such as sliding and hopping (**Fig. 7B**). This contrasts with the *E. coli* holoenzyme, which primarily explores the genome through three-dimensional diffusion ^51,52^. This difference may be functionally related to the distinct cellular morphologies: *E. coli* cell is a 0.5 × 1-3 µm rod with a compact nucleoid, while *S. africana* is 0.25-0.3 µm thick and 15-30 µm long ^26^. Given that the spirochetal nucleoid stretches along the entire length of the bacterium ^53^, such a highly anisotropic environment may render three-dimensional diffusion inefficient, necessitating one-dimensional facilitated diffusion for efficient genome exploration.

Our study of the spirochetal transcriptional apparatus reveals significant phylum-level functional diversity of RNAPs within the Pseudomonadati kingdom. These findings underscore the importance of thorough biochemical and structural characterization of diverse RNAPs and lay the foundation for understanding transcription regulation in pathogenic spirochetes such as *Treponema*, *Borrelia*, and *Leptospira*.

## Methods

### Reagents

NTPs from Jena Bioscience, Bioline Reagents and Thermofisher were used interchangeably in *in vitro* transcription assays. DFHBI-1T (CAS Number: 1539318-36-9) was from Jena Bioscience and TargetMol. Other reagents were purchased from Merck and Fisher Scientific.

### Molecular phylogenetic analysis

Protein sequences of RNAP β, β’, α subunits and NusA transcription elongation factor were fetched from the KEGG database ^54^. 2-5 divergent sequences per phylum were selected (**Supplementary Table 1**) to achieve balanced representation of different bacterial phyla. Protein sequences were concatenated, aligned with Muscle (default settings) ^55^ and conserved blocks of sequences were extracted using Gblocks (default settings) ^56^. A molecular phylogenetic tree was constructed using MrBayes software ^57^ using the empirical Jones-Taylor-Thornton amino acid substitution matrix^58^, the rate heterogeneity across sites was modeled with 8 gamma categories ^59^. The analysis was run for 200,000 generations, trees were sampled every 1,000 generations, the first 50 trees (50,000 generations) were discarded, and the consensus tree was built from the remaining 150 trees. Seaview ^60^ and Ugene ^61^ software packages were used as graphical user interfaces to run Muscle, Gblocks, MrBayes and visualize molecular phylogenetic trees. For indel analysis, the correspondence (likely homology) between positionally aligned lineage-specific inserts (Si1, Si2 and Si3) was estimated based on experimentally determined structures (where available) or AlphaFold ^62^ models of individual RNAP subunits (precomputed ^63^ or generated using the AlphaFold Server (https://alphafoldserver.com/) ^64^).

### Plasmids for protein expression in *E. coli*

Plasmids employed for protein expression are presented in **Supplementary Table 3**. Wild-type and variant *S. africana* and *E. coli* RNAPs were produced from polycistronic vectors encoding αββ’ω subunits with polyhistidine tags at the C-terminus of β’, the N-terminus of β or both. Wild-type and variant *S. africana* σ factors, CarD, DksA, *E. coli* σ factor and DksA were produced from plasmids encoding gene fusions with N-terminal polyhistidine tags followed by a TEV, ULP1 (SUMO domain) or thrombin protease site.

### Plasmids used as transcription templates

All transcription template plasmids (**Supplementary Table 4**) were derivatives of pOP004 and contained the identical ∼2.4 kb pOA-RQ backbone (containing the ColE1 origin and encoding β-lactamase) and a ∼1 kb insert between *Sfi*I sites. The insert contained a transcription unit consisting of a promoter and a 0.8 kb transcribed region ending with a strong hairpin-dependent terminator. The transcribed region encoded a Broccoli fluorogenic light up aptamer ^65^ embedded in a tRNA scaffold just upstream of the terminator. The upstream 0.2 kb of the insert was unique for each plasmid and contained an *Xba*I-*Nhe*I module encompassing a promoter with flanking genomic sequences (40 bp upstream of the −35 element and 20 bp downstream of the transcription start site). The downstream 0.8 kb of the insert were the same for all plasmids except for pJK006 and pJK004 contained a featureless *Nhe*I-*Mfe*I module, a *Mfe*I-*BsrG*I module encoding a Broccoli aptamer, followed by a *BsrG*I-*Xho*I module encoding the *E. coli hisT* terminator. In the pJK006 and pJK004 plasmids, phage λ Rho utilization sites were inserted near the upstream edge of the *Nhe*I-*Mfe*I module.

Plasmids stocks (1-2 µM) were produced from 0.5-1 l of LB culture using NucleoBond Xtra Maxi kit (Macherey-Nagel). 500 nM stocks of linearized plasmids were prepared by digestion with *Xho*I restriction enzyme in the Thermofisher Buffer R (10mM Tris-HCl pH 8.5 at 37°C, 10 mM MgCl_2_, 100 mM KCl, 0.1 mg/ml BSA) and used directly in *in vitro* transcription assays.

### Protein production in *E. coli*

Individual components of *S. africana* and *E. coli* transcription systems were expressed in *E. coli. E. coli* Xjb(DE3), BL21(DE3) and T7 Express lysY/Iq (New England Biolabs) stains were used interchangeably unless indicated otherwise. Cultivation was performed in 4 l of LB medium in flasks or a fermenter. Cells were harvested by centrifugation at 7,000 × g for 20 min at 4°C. Cultivations yielded from 15 g (5 h induction) to 20-40 g (overnight induction) of cells. *E. coli* proteins were produced using 5 h induction with IPTG at 30°C. *S. africana* proteins other than σ factors were produced using the overnight express autoinduction protocol ^66^ at 37°C (≥18 h). *S. africana* σ factor was produced in *E. coli* T7 Express *lys*Y/I^q^ cells. Cells were pre-grown to OD_600_ 0.6 and induced with 1 mM IPTG overnight at 25°C.

### Purification of proteins produced in *E. coli*

Wild-type and variant RNAPs and σ factors were purified by combination of Ni^2+^-affinity, heparin and anion exchange chromatography ^67^. Cells (20-40 g) were resuspended in Lysis Buffer (50 mM Tris-HCl pH 7.9, 5% (v/v) glycerol, 500 mM NaCl, 1 mM β-mercaptoethanol) supplemented with EDTA-free protease inhibitor cocktail (Pierce or Roche). At least 30 ml of lysis buffer was used per 5 g of cells. When using *E. coli* BL21(DE3) or T7 Express lysY/Iq cells, the lysis buffer was supplemented with 1 mg/ml of lysozyme and the cells were incubated on ice for 1 h prior to disruption by sonication, whereas *E. coli* Xjb(DE3) cells were disrupted immediately following the resuspension in lysis buffer without lysozyme. Lysates were clarified by centrifugation at 40,000 × g for 30 min at 4°C. Clarified lysates were loaded onto 3 gravity columns each containing 1 ml of WorkBeads NiMAC (Bio-Works) chromatography matrix.

Columns were washed with 10 ml of lysis buffer, and the target protein was eluted with a step gradient of imidazole (50, 100 and 200 mM, 10 ml each passed sequentially through the 3 columns) in Lysis Buffer. Fractions containing the target proteins were diluted three times with buffer A (50 mM Tris-HCl pH 7.9, 5% (v/v) glycerol, 1 mM β-mercaptoethanol, 0.1 mM EDTA) and loaded onto a 5 ml heparin column (HiTrap® Heparin HP, Cytiva) equilibrated with 10% buffer B (50 mM Tris-HCl pH 7.9, 1.5 M NaCl, 5% (v/v) glycerol, 1 mM β-mercaptoethanol, 0.1 mM EDTA), washed with 15-30 ml of 10% buffer B, and the target protein was eluted with a 100 ml gradient of buffer B (10-100%). Fractions containing the target protein were pooled, diluted 2-3 times with buffer A and loaded onto a 6 ml ResourceQ column (Cytiva) equilibrated with 10% buffer B, washed with 15-30 ml of 10% buffer B, and the target protein was eluted with a 100 ml gradient of buffer B (10-100%). Fractions containing the target protein were pooled, concentrated using ultrafiltration concentrators, dialyzed against storage buffer (50% (v/v) glycerol, 20 mM Tris-HCl pH 7.9, 150 mM NaCl, 0.1 mM EDTA, 0.1 mM DTT) and stored at −20°C (short term) or −80°C (long term). For target proteins with cleavable N-terminal polyhistidine tag, removal of the tag was performed either after elution from WorkBeads NiMAC or after heparin chromatography by overnight digestion with in-house produced TEV (His_6_-TEV-gene constructs) or in-house produced ULP1 protease (His_6_-SUMO-gene constructs). Prior to digestion with proteases, the NaCl concentration was reduced to 0.3 M by dilution with buffer A. *S. africana* CarD, DksA and *E. coli* DksA were purified using capture on WorkBeads NiMAC as described above followed by cleavage of the polyhistidine tag and size exclusion chromatography on a 120 ml HiPrep™ Sephacryl™ S-200 HR 16/60 column (Cytiva) in 33 or 50% buffer B.

### Production of *Sfc* RNAP *via* recombinant baculovirus in insect cells

Plasmid pFL-AC_pUCDM-BZ was created from the donor plasmid pUCDM (with *S. africana rpoB* and *rpoZ* genes encoding β and ω subunits, respectively) and the acceptor plasmid pFL (with *S. africana rpoA* and *rpoC* genes encoding α and β′ subunits, with β′ fused to a C-terminal His_6_-tag) *via* Cre-LoxP recombination ^68^. The resulting pFL-AC_pUCDM-BZ plasmid was introduced into DH10MultiBac *E. coli* cells to produce recombinant bacmid. The bacmid was transfected into Sf9 cells to produce generations V1 and V2 of recombinant baculovirus particles. 400 ml Hi5 cell culture at 0.8 x 10^6^ cells/ml were infected with 1 ml V2 virus. Cells were further cultured at 27 °C with shaking (80 rpm), split after 24 h and, after an additional 48 h incubation, harvested by centrifugation.

All purification steps were conducted at 4 °C. The cell pellet was resuspended in 40 ml lysis buffer (50 mM Tris-HCl, pH 6.9, 500 mM NaCl, 5 % (v/v) glycerol), supplemented with cOmplete^TM^ EDTA-free protease inhibitor (Sigma-Aldrich), 1 mg/ml DNase, 1 mM β-mercaptoethanol and 0.2 % (v/v) Tween 20, and disrupted by sonication. The lysate was cleared by centrifugation, and the supernatant was loaded onto a Ni^2+^-NTA HisTrap column (Cytiva) pre-equilibrated with lysis buffer. Proteins were eluted using a linear gradient (20 - 250 mM) of imidazole in lysis buffer. Fractions containing *S. africana* RNAP were identified by SDS PAGE, pooled and diluted 1:4 with buffer A (50 mM Tris-HCl, pH 6.9, 5 % (v/v) glycerol, 1 mM β-mercaptoethanol, 0.1 mM EDTA). *S. africana* RNAP was further purified using HiTrap Heparin HP and HiTrap MonoQ HP columns (Cytiva) with analogous dilution in between and using gradients to buffer A plus 1.5 M NaCl. Fractions containing *S. africana* RNAP were pooled, and the target was further purified by size exclusion chromatography (SEC) on a HiLoad Superdex 200 Increase 16/600 column (Cytiva) in storage buffer (50 mM Tris-HCl, pH 6.9, 300 mM NaCl, 0.1 mM EDTA, 2 mM DTT, 10 % (v/v) glycerol). Fractions containing *S. africana* RNAP were concentrated to approximately 8 mg/ml, aliquoted, flash-frozen in liquid nitrogen, and stored at −70 °C.

*S. africana* σ^70^ holoenzyme was prepared by mixing 10 µM (yielding monomeric complexes) or 20 µM (yielding dimeric complexes) RNAP with a threefold molar excess of σ^70^ and incubation for 10 min at 32 °C. Template (tDNA) and non-template DNA (ntDNA) were mixed at equimolar concentrations in binding buffer (20 mM Tris-HCl, pH 7.9, 120 mM KOAc, 5 mM Mg(OAc)_2_, 10 µM ZnCl_2_, 2 mM DTT) and annealed by heating to 95 °C for 5 min and subsequent cooling to 25 °C at 1 °C/min. The annealed scaffolds were incubated with *S. africana* σ^70^ holoenzyme in a 1.3:1 molar ratio for 10 min on ice, followed by an additional incubation for 10 min at 32 °C. The mixture was subjected to SEC on a Superdex 200 Increase 3.2/300 column in binding buffer. Fractions containing *S. africana* σ^70^ promoter complexes were identified by SDS PAGE and urea PAGE analysis, pooled and concentrated to 5-6 mg/ml by using a 100 kDa cutoff Amicon Ultra Centrifugal Filter (Sigma-Aldrich).

*S. africana* CarD-modified σ^70^ promoter complexes were prepared by mixing 20 µM RNAP with σ^70^ and, subsequently, CarD, each at threefold molar excess, with incubation for 10 min at 32 °C after each step. Equimolar tDNA and ntDNA were annealed as above, mixed with *S. africana* CarD/holoenzyme mix in a 1.3:1 molar ratio and incubated for 10 min on ice, followed by incubation for 10 min at 32 °C. The mixture was subjected to SEC in binding buffer as above. Fractions containing CarD-modified σ^70^ promoter complexes were pooled and concentrated to 5-6 mg/ml as above.

### Fluorogenic light-up aptamer *in vitro* transcription assays

FLAP assays were performed largely as described previously ^28^. Reactions were initiated by mixing 35 µl of holoenzyme-DNA solution with 35 µl of NTP solution. Both solutions contained the transcription buffer and were pre-warmed at 37°C. Within 20 s, 55 µl of the reaction mix was transferred to the ultra-micro fluorometer cuvette (Hellma 105.251-QS) pre-warmed at 37°C, the cuvette was placed in the fluorometer equipped with a thermostated cuvette holder at 37°C, and fluorescence was recorded in a time-drive mode for 10 min. The reaction mixture contained 10 µM of (5Z)-5-[(3,5-difluoro-4-hydroxyphenyl)methylene]-3,5-dihydro-2-methyl-3-(2,2,2-trifluoroethyl)-4H-imidazol-4-one (DFHBI-1T), 1 mM each of ATP, CTP, GTP and UTP, 50 nM of linearized plasmid DNA template, 100 nM yeast inorganic pyrophosphatase, 1 µM RNAP, 4 µM σ.

Transcription buffers contributed 10 mM MgCl_2_, 80 mM KCl, 0.1 mM EDTA, 0.1 mM DTT and 40 mM of zwitterionic buffer (HEPBS-KOH pH 9.0, HEPBS-KOH pH 8.5, HEPPS-KOH pH 8.0, HEPES-KOH pH 7.5; all pH values at 25°C) to the reaction mixture. Taking into account the contributions of the storage buffer (from proteins), buffer R (from DNA templates) and NTPs (disodium salts), the final reaction mixture contained 11 mM MgCl_2_, 110-120 mM KCl, 25 mM NaCl, 3 mM Tris, 5% (v/v) glycerol, 1% (v/v) DMSO, and 0.01 mg/ml BSA.

Fluorescence time curves were highly reproducible within the single experiment and across independent experiments performed days apart with the same batch of RNAP. However, the absolute activities of different batches of RNAP differed by as much as twofold and decreased upon storage on the timescale of weeks. All major effects reported in this study were confirmed by the direct comparison within the same experiment. The reported absolute activities were adjusted based on activity of the P-gre promoter that was measured in each experiment. A median activity of the P-gre promoter with fresh batches of RNAP was used as a reference.

In a subset of experiments, we employed an end-point-only modification of the FLAP assay to facilitate rapid acquisition of data. Reactions were initiated by mixing 20 µl of holoenzyme-DNA solution with 20 µl of NTP solution. Both solutions were pre-warmed at 37°C and did not contain DFHBI-1T and 1% DMSO. The reactions were incubated for 10 min in a thermoblock at 37°C and quenched by 20 µl of solution containing 45 mM EDTA (15 mM final) and 30 µM DFHBI-1T (10 µM final). For zero samples, holoenzyme-DNA solution was mixed with quench solution before adding NTP solution. 55 µl of the quenched reaction mix were transferred to the fluorometer cuvette, the cuvette was placed in the fluorometer equipped with a thermostated cuvette holder at 25°C, and the fluorescence spectrum was recorded (490-600 nm, excitation at 472 nm). Fluorescence at 507 nm corrected for the fluorescence of the zero samples and multiplied by 1.5 to account for the dilution during quenching was then used as a measure of transcription activity.

### Denaturing PAGE analysis of RNA products

Transcription reactions were set up as described in the FLAP assay section. A 20 µl aliquot was withdrawn after 10 min and quenched by 60 µl of Gel Loading Buffer (94% formamide, 20 mM Li_4_-EDTA and 0.2% OrangeG). RNAs (1.5 µl) were separated on 6% 10 × 12 cm denaturing TBE-urea polyacrylamide gels (30 min at 70 V followed by 75 min at 180 V) in 1×TBE buffer, stained with SYBR-Gold for 30 min in 1×TBE buffer, and visualized with a Sapphire FL Biomolecular Imager (Azure Biosystems). Gel images were processed using the ImageJ software ^69^.

### Analysis of RNA products using Qubit fluorometer

Transcription reactions (70 µl, in order to use the same pipetting volumes as in the FLAP assays) were set up as described in the FLAP assay section except that DFHBI-1T was omitted. Reactions were quenched after 10 min by withdrawing a 40 µl aliquot and mixing it with 10 µl of 0.5 M EDTA (100 mM final). For zero samples, EDTA was added to the holoenzyme-DNA solution before mixing with the NTP solution. 2-20 µl of quenched reactions were mixed with Qubit Broad Range RNA assay reagent (Thermofisher) to achieve the final volume of 200 µl. The apparent RNA concentration was then determined by measuring the sample fluorescence using the Qubit instrument (version 4, Thermofisher) calibrated using RNA standards. The actual RNA concentration in the sample was then estimated by subtracting the zero sample readings (∼25 nM).

### CryoEM grid preparation, data acquisition and processing

Samples were supplemented with 0.15 % (w/v) n-octylglucoside. A 3.8 µl aliquot of the sample was applied to glow-discharged Quantifoil R 1.2/1.3 Cu grids at 10 °C and 100 % humidity, and blotted using a FEI Vitrobot Mark IV with a blot force setting of −10, a waiting time of 4 s and a blotting time of 4 s. The grids were then plunged into liquid ethane, cooled by liquid nitrogen, and stored in liquid nitrogen until data collection.

Images were collected at a nominal magnification of 96,000x in counting mode, using a FEI Titan Krios G3i transmission electron microscope (TEM) at 300 kV, equipped with a Falcon 3EC camera, using the EPU software (ThermoFisher Scientific) with a calibrated pixel size of 0.832 Å. A total electron dose of 40 e^−^/Å^2^ was accumulated over an exposure time of 40.57 s. Movie alignment was done with Patch Motion correction, followed by contrast transfer function (CTF) estimation with PatchCTF within cryoSPARC ^70^.

For cryoEM image processing, all steps were performed using cryoSPARC ^70^. For monomeric *S. africana* σ^70^ promoter complexes, 6,990 micrographs were collected (**Supplementary Fig. 7**). Micrographs showing defects in the Thon rings due to excessive drift, ice contamination or astigmatism were discarded. An initial model for reference-based particle selection from micrographs was created using class averaging of manually selected particles. In the initial analysis, particle images were extracted by using a box size of 440, binned to 110. All raw particles were automatically selected and submitted for 2D classification to remove poor-quality particles. After 2D classification, 1,925,988 particles were retained for further analysis. *Ab initio* reconstruction was conducted to generate an initial 3D reference for 3D heterogeneous refinement, using a small subset of particles. The dataset was iteratively classified into three populations, and two bad populations were discarded. Selected particles were re-extracted from the good population (849,917 particles) with a box of 220 and 3D classified again to clean the dataset further. Finally, selected particle images were re-extracted with a box of 280 (1.3 Å/px) and subjected to local refinement using a generously enlarged soft-mask.

After per-particle CTF correction, non-uniform refinement ^71^ was applied to generate the final cryoEM maps with an estimated average resolution of 3.0-3.4 Å according to the gold-standard Fourier shell correlation cutoff of 0.143 ^72^ (**Supplementary Figs. 7-8**). CryoEM images of dimeric *S. africana* σ^70^ promoter complexes (**Supplementary Figs. 9-10**) and of CarD-modified σ^70^ promoter complexes (**Supplementary Figs. 11-12**) were processed in an analogous fashion.

### Model building, refinement and analysis

AlphaFold3 models ^64^ of the individual proteins were computed and manually placed into the cryoEM reconstructions, adjusted by rigid body fitting and segmental real-space refinement using Coot (version 1.11.1) ^73,74^. The structural models were refined by iterative rounds of real space refinement in PHENIX (version 1.21.2-5419) ^75,76^ and manual adjustment in Coot. The structural models were evaluated with MolProbity (version 4.5.1) ^77^. Structure figures were prepared with PyMOL (version 2.4.0, Schrödinger) and ChimeraX ^78^.

### EMSA assays

DNA fragments were produced by preparative PCR (0.5-1 ml total volume). Plasmid pVN001, DY782-GGAGAGACAACTTAAAGAGACATTCTG and DY682-TCGTAGATTGTTATTCAACT primers were used to prepare a 163 bp DNA fragment with the P-gre promoter: GGAGAGACAACTTAAAGAGACATTCTGCATCATGATATTGTTATTATGCTTCCTGCGGCATATACCCTGCCCATTCTGTGCGCATACCATCCCTGCCTGCGTTGACGAAACTCGTTAACATGTTATAATACAAGCGTTACCATAGTTGAATAACAATCTACGA. Plasmid pVL026, DY782-GGAGAGACAACTTAAAGAGACATTCTG and DY682-TCCTCTGGTTCCTCGTGTCC primers were used to prepare a 163 bp promoter-less DNA fragment:

GGAGAGACAACTTAAAGAGACATTCTGCATCATGATATTGTTATTATGCTTCCTGCGGCATATACCCTGCCCATTCTGTGCGCATACCATCCCTGCCTGCGCTGCAGGCACTGTTTACCGCAGTGCTGCACAAGCATCGAGAGGGACACGAGGAACCAGAGGA. DNA fragments were purified using the NucleoSpin Gel and PCR Clean-up kit (Macherey-Nagel GmbH). Reactions were prepared by mixing 1 µl of RNAP and 1 µl of σ in storage buffer with 8 µl of DNA fragment in 1.25 × reaction buffer, resulting in 10 µl samples containing 25 nM of DNA fragment, desired concentrations of RNAP and σ, 40 mM HEPBS-KOH pH 9.0, 80 mM KCl, 15% glycerol, 0.1 mM EDTA and 0.1 mM DTT. Reactions were incubated for 20 min at 30°C, mixed with 10 µl of loading solution (20% glycerol and 0.4% OrangeG), and 2 µl was loaded onto a 4% 10 × 12 cm 1×TBE polyacrylamide gel. Electrophoresis was performed for 35 min at 50 V, followed by 25 min at 100 V. DNA bands were visualized with an Odyssey Infrared Imager using 700 nm channel (Li-Cor Biosciences). Gel images were processed using the ImageJ software^69^.

## Data availability

CryoEM reconstructions have been deposited in the Electron Microscopy Data Bank (https://www.ebi.ac.uk/pdbe/emdb) under accession codes: 56405, 56406, 56407, 56408 for monomeric RPo-P-gre (state I to IV), 56409 for dimeric RPo-P-gre, 56410 for CarD-RPo-P-gre as well as 56411 for CarD-RPo-P-rrna. Structure coordinates have been deposited in the RCSB Protein Data Bank (https://www.rcsb.org) under accession codes: 9TXX, 9TXY, 9TXZ, 9TX0 for monomeric RPo-P-gre (state I to IV), 9TX1 for dimeric RPo-P-gre, 9TX2 for CarD-RPo-P-gre as well as 9TX3 for CarD-RPo-P-rrna. RNA-seq data have been deposited in the GEO database (https://www.ncbi.nlm.nih.gov/geo/query/acc.cgi?acc=GSE305863)

CLMS data have been deposited in jPOST (https://jpostdb.org/) with accession code JPST004380 (PXID PXD074046)^79^

All other data are contained in the manuscript or the Supplementary Information. Supplementary Data File is provided in this paper.

## Acknowledgements

Work in the authors’ laboratories was supported by grants from the Academy of Finland (341962 to G.A.B.), the Sigrid Jusélius Foundation (to G.A.B.) and the Deutsche Forschungsgemeinschaft (WA 1126/11-1 to M.C.W.). We thank Irina Artsimovitch, Ohio State University, for critically reading the manuscript. We acknowledge the assistance of the Core Facility BioSupraMol supported by the German Research Foundation (Deutsche Forschungsgemeinschaft). We are grateful to Jörg Bürger, Freie Universität Berlin, Research Center of Electron Microscopy and Core Facility BioSupraMol, for help with cryoEM sample preparation. The study utilized protein expression and purification infrastructure provided by Turku Protein Core. Transcriptome sequencing was performed by the Finnish Functional Genomics Centre, the University of Turku and Åbo Akademi, supported by the Biocenter Finland. The authors wish to acknowledge CSC – IT Center for Science, Finland, for computational resources. Mass spectrometry analyses were performed at the Turku Proteomics Facility supported by Biocenter Finland. We thank Verneri Nissilä, Iida Payne, Oskari Puro and Karina Šapovalovaitė for their assistance with assay development and experimental condition screening during the initial phases of this study.

## Author contributions

V.K.T. most FLAP assays, RNA isolation from spirochetes. T.W. cryoEM analyses, sample preparation, model building. T.H. cryoEM analyses and data processing. J.J.M. reconstitution of spirochetal transcription *in vitro*. J.K. nanopore sequencing, transcriptome analysis. P.J.T. EMSA experiments. V.L. FLAP assays. B.L. cryoEM analyses, model building and refinement. Protein production: T.W., J.J.M., S.P., V.K.T., V.L., J.K. Plasmids construction: T.W., J.J.M., V.L., J.K. Large scale DNA production: V.L., J.K., V.K.T. Crosslinking mass-spectrometry: V.K.T, S.P., J.J.M. Cultivation of spirochetes: V.K.T, S.P., J.K. All authors contributed to the analysis of the data and the interpretation of the results. M.C.W. and G.A.B. conceived, designed, supervised the study and wrote the manuscript with contributions from the other authors.

## Conflict of interest

The authors declare no conflict of interest.

## Supplementary Information

### Supplementary methods

#### S. africana cultivation

*S. africana* was routinely cultivated in an alkali medium (pH 9.2-9.5) modified from the DSMZ alkaliphilic spirochaete medium #700. The medium was buffered by a mixture of Na_2_CO_3_ (10 g/l) and NaHCO_3_ (15 g/l), contained NaCl (50 g/l) to maintain high-salinity ^1^, maltose (5 g/l) as a major carbon and energy source and Na_2_S (0.5 g/l) and cysteine hydrochloride (0.5 g/l) as reducing agents. The medium also contained NH_4_Cl (0.5 g/l), K_2_HPO_4_ (0.5 g/l), yeast extract (0.5 g/l), 10 ml/l of vitamin solution, 1 ml/l of trace element solution and 1 mg/l of resazurin as redox indicator. For cultivation at pH 8.5, NaHCO_3_ was increased to 25 g/l and Na_2_CO_3_ was omitted. For monitoring the survival at pH 7.5, HEPES-Na (HEPES 12 g/l, NaOH 1 g/l) was used as a buffering agent, Na_2_CO_3_ and NaHCO_3_ were omitted, and NaCl was increased to 90 g/l.

A continuous culture was grown in completely filled 6.5 ml soda-lime glass tubes with polypropylene screw caps and thermoplastic elastomer seal (or up to 1 l screw caps bottles when desired) without shaking at 25°C, pH 9.2-9.5 and transferred every 3 days (OD_600_ 0.3-0.5; 25 µl to 6.5 ml). Alternatively, 3-day old cultures were supplemented with glycerol (16% final), stored at −80°C and revived when desired. On day 4 cultures reached saturating OD_600_ ∼0.8, cells maintained spiral shape and motility, and the culture remained evenly turbid. On day 5 culture health deteriorated, a large fraction of cells sedimented to the bottom of the tube and morphed into spherical bodies. When grown at 30°C, cultures reached saturation on day 3 and the viability deteriorated on day 4. When grown at 25°C, pH 8.5, cultures reached saturation on day 7 and remained healthy on day 12 after which the viability rapidly deteriorated. When 3-day old cultures pre-grown at pH 9.2-9.5 were pelleted and transferred to pH 7.5 medium, cells remained spiral and motile on day 2 followed by the massive die out on days 3-4, but a small number of spiral and motile cells were observed up to at least day 7.

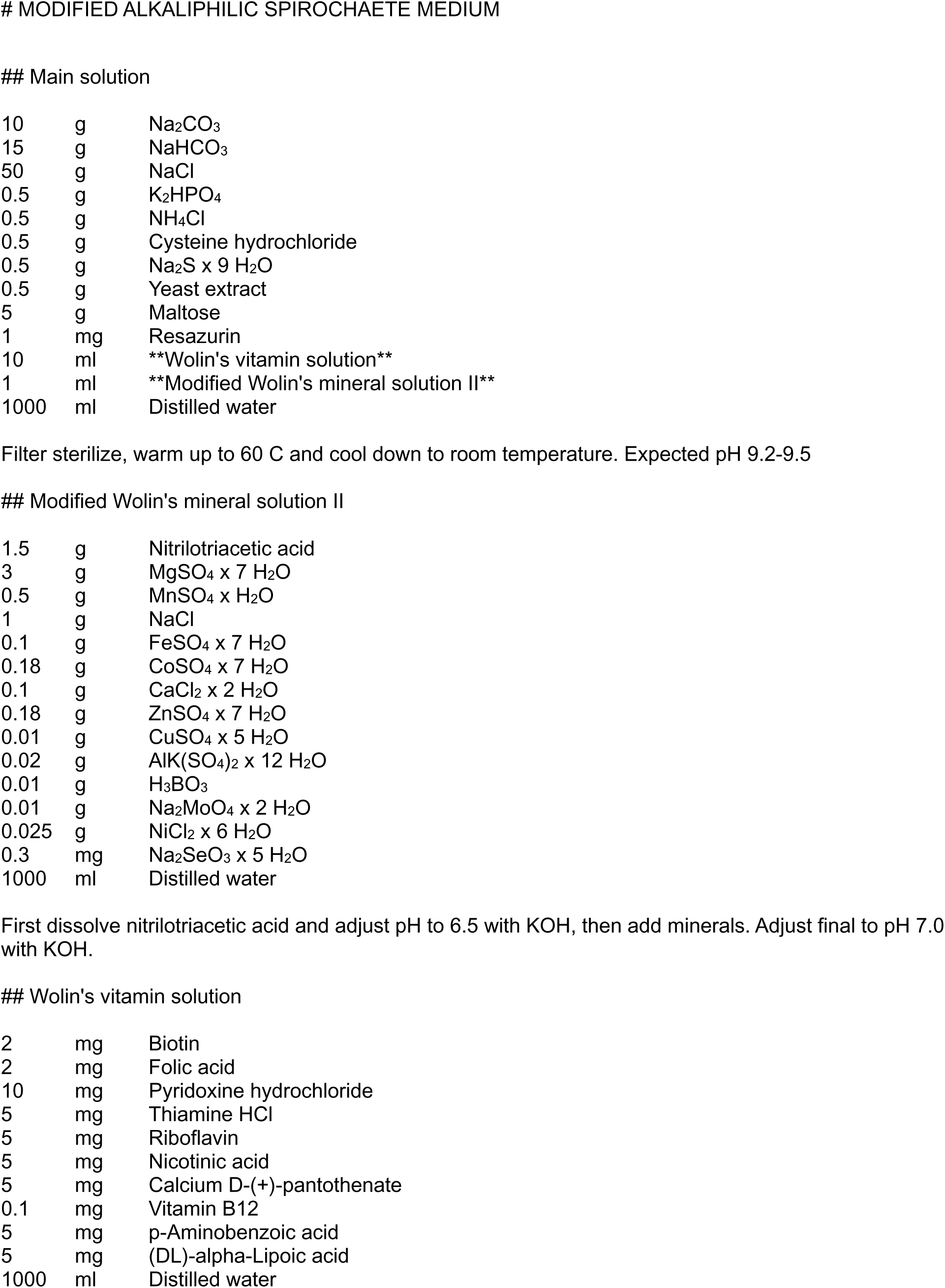

### Main solution
|  |  |  |
| --- | --- | --- |
| 10 | g | Na<sub>2</sub>CO<sub>3</sub> |
| 15 | g | NaHCO<sub>3</sub> |
| 50 | g | NaCl |
| 0.5 | g | K<sub>2</sub>HPO<sub>4</sub> |
| 0.5 | g | NH<sub>4</sub>Cl |
| 0.5 | g | Cysteine hydrochloride |
| 0.5 | g | Na<sub>2</sub>S x 9 H<sub>2</sub>O |
| 0.5 | g | Yeast extract |
| 5 | g | Maltose |
| 1 | mg | Resazurin |
| 10 | ml | **Wolin's vitamin solution** |
| 1 | ml | **Modified Wolin's mineral solution II** |
| 1000 | ml | Distilled water |
| Modified Wolin's mineral solution II |  |  |
| 1.5 | g | Nitrilotriacetic acid |
| 3 | g | MgSO<sub>4</sub> x 7 H<sub>2</sub>O |
| 0.5 | g | MnSO<sub>4</sub> x H<sub>2</sub>O |
| 1 | g | NaCl |
| 0.1 | g | FeSO<sub>4</sub> x 7 H<sub>2</sub>O |
| 0.18 | g | CoSO<sub>4</sub> x 7 H<sub>2</sub>O |
| 0.1 | g | CaCl<sub>2</sub> x 2 H<sub>2</sub>O |
| 0.18 | g | ZnSO<sub>4</sub> x 7 H<sub>2</sub>O |
| 0.01 | g | CuSO<sub>4</sub> x 5 H<sub>2</sub>O |
| 0.02 | g | AlK(SO<sub>4</sub>)<sub>2</sub> x 12 H<sub>2</sub>O |
| 0.01 | g | H<sub>3</sub>BO<sub>3</sub> |
| 0.01 | g | Na<sub>2</sub>MoO<sub>4</sub> x 2 H<sub>2</sub>O |
| 0.025 | g | NiCl<sub>2</sub> x 6 H<sub>2</sub>O |
| 0.3 | mg | Na<sub>2</sub>SeO<sub>3</sub> x 5 H<sub>2</sub>O |
| 1000 | ml | Distilled water |
| Wolin's vitamin solution |  |  |
| 2 | mg | Biotin |
| 2 | mg | Folic acid |
| 10 | mg | Pyridoxine hydrochloride |
| 5 | mg | Thiamine HCl |
| 5 | mg | Riboflavin |
| 5 | mg | Nicotinic acid |
| 5 | mg | Calcium D-(+)-pantothenate |
| 0.1 | mg | Vitamin B12 |
| 5 | mg | p-Aminobenzoic acid |
| 5 | mg | (DL)-alpha-Lipoic acid |
| 1000 | ml | Distilled water |

#### Purification of *S. africana* RNAP from *S. africana* cells

Native *S. africana* RNAP was purified from *S. africana* cells by a combination of heparin, size-exclusion and anion exchange chromatography. *S. africana* cultures were grown for 3-4 days at 25-30°C until the early saturation at OD_600_ ∼0.8 (6.5 ml inoculum per 1 l bottle of standard medium with pH 9.2-9.5). *S. africana* cells were harvested by centrifugation at 10,000 × g for 30 min at 4°C (∼15 g from 6 l of culture, stored at −80°C), resuspended in 10% Buffer B supplemented with EDTA-free protease inhibitor cocktail (Pierce or Roche) and disrupted by a mild sonication. At least 30 ml of lysis buffer was used per 5 g of cells.

Lysates were clarified by centrifugation at 40-60,000 × g for 0.5-1 h at 4°C and loaded onto a 5 ml HiTrap® Heparin HP column (Cytiva) equilibrated with 10% Buffer B. The column was washed with 15 ml of 10% Buffer B and proteins were eluted with a steep gradient of Buffer B (30 ml, 10-100%). All protein-containing fractions were pooled, concentrated to 2 ml using a Amicon Ultra-15 (15 ml) centrifugal filter unit with Ultracel-3 (3 kDa cutoff) membrane and loaded onto a 120 ml HiPrep™ Sephacryl™ S-400 HR 16/60 gel filtration column (Cytiva) equilibrated with 10% Buffer B. RNAP eluted from the column between 70 and 80 ml. Fractions containing RNAP were pooled, concentrated using ultrafiltration concentrators, dialyzed against the storage buffer and stored at −20°C (short term) or −80°C (long term). Alternatively, RNAP-containing fractions were loaded onto a 6 ml ResourceQ column (Cytiva) equilibrated with 10% Buffer B, washed with 15-30 ml of 10% buffer B, and RNAP was eluted with a 100 ml gradient of buffer B (10-100%) followed by concentration and dialysis.

#### Total RNA isolation from *S. africana*

Total cell RNA was isolated using TRIzol® Max™ Bacterial RNA Isolation Kit (Thermofisher). Cells from 6.5 ml culture were harvested by centrifugation at 4,000 × g for 10 min at 4°C and resuspended in 200 µl of Max Bacterial Enhancement Reagent preheated at 95°C. The sample was then dissolved in 1 ml TRIzol and extracted twice with 200 µl of cold chloroform. RNA was then precipitated from the aqueous phase by addition of 500 µl of cold isopropanol. The RNA pellet was washed with 1 ml of 75% ethanol and dissolved in 50 μl of RNase-free water. The sample was then treated with DNase (TURBO DNA-free™ Kit, Thermofisher). Briefly, the sample was supplemented with DNase buffer, DNase, diluted to 100 µl, and incubated for 20 min at 37°C. DNase and divalent cations were then removed by the proprietary DNase removal matrix supplied with the kit. RNA was then additionally cleaned up using Qiagen RNAeasy mini kit using Clean-up protocol. Briefly, the RNA sample was supplemented with 350 µl of high-salt proprietary buffer containing guanidine thiocyanate (Buffer RLT), 950 µl of 99% ethanol (up from the default 250 µl of ethanol to facilitate binding of small RNAs) and applied to the silica-based spin column. The column was washed with the low-salt proprietary buffer (Buffer RPE) containing 75% ethanol, and RNA was eluted with 50 µl of RNase-free water.

#### Transcriptome sequencing and analysis

Total RNA was isolated from a 3-day-old *S. africana* culture grown at pH 9.2-9.5 until early saturation (OD_600_ ∼0.8) at 30 °C. Total RNA (100 ng) was converted into a dual-indexed library using Illumina® Stranded Total RNA Prep, Ligation with Ribo-Zero Plus kit and sequenced with Illumina NovaSeq 6000 SP v1.5 (2 × 100 bp). The library preparation and RNA sequencing were performed by the Finnish Functional Genomics Centre Facility (FFGC). The RNA sample and library quality were monitored using an Agilent Bioanalyzer 2100 instrument and software (Agilent). The read quality was inspected using FastQC.

The automated transcriptome assembly was conducted using Rockhopper v2.03 ^2^ that aligned raw reads (FASTQ format) to the annotated *S. africana* DSM8902 genome ^3^, produced coverage traces (WIG format) and a table containing raw read counts and expression values (analogues of FPKM but normalized by the upper quartile of gene expression, excluding genes with zero expression) for annotated genome features (**Supplementary Data File**). Transcriptome coverage was explored using Integrative Genomics Viewer ^4^. For figure preparation, strand-specific transcriptome coverage of selected regions of the genome was extracted from WIG files using a python script and plotted using Origin 2015 software (Origin Labs).

As an alternative analysis method, raw reads (FASTQ format) were mapped to the *S. africana* genome using Bowtie2 v2.5.3 ^5^. The resulting alignment in SAM format was sorted and indexed using SAMtools v1.19 ^6^. Reads mapping to the genome features (Gene Transfer Format, GTF) were then counted using HTSeq v2.0.3 ^7^. The HTSeq read counts were normalized using the transcripts per million (TPM) method (**Supplementary Data File**).

#### Analysis of *in vitro* transcription products using Oxford nanopore sequencing

To estimate transcription across the entire linearized transcription template (P-gre template was used as an example), *in vitro* transcription was performed as in the FLAP assays, but the reaction volume was scaled up to 0.5 ml and DFHBI-1T was omitted.

Product RNAs were purified using the RNA easy kit (Qiagen) and polyadenylated with *E. coli* PAP (poly-A polymerase, NEB) to enable ligation of the sequencing adapters to the 3’ ends of the RNAs. RNAs were purified using Agencourt RNAClean XP beads (Beckman Coulter), the sequencing library was prepared using the Direct RNA sequencing kit (Oxford nanopore) and sequenced using a MinION Mk1D instrument (Oxford nanopore). Reads were aligned to the linearized P-gre transcription template using MinKNOW, alignment files were split according to the strand using SAMtools^6^, loaded into Ugene^8^, strand-specific coverage was exported and ultimately plotted using Origin 2015 software.

In the transcription start site mapping experiment, seven *in vitro* transcription reactions were performed individually, each using a dedicated linearized plasmid template (six plasmids from the main set and the parent pOP004 plasmid featuring P-gre promoter with the altered initially transcribed region). One set of reactions was performed using *S. africana* holoenzyme and another set using *E. coli* holoenzyme. After quenching with EDTA, the product RNAs in each set were pooled together, purified using the RNA easy kit and polyadenylated with *E. coli* PAP. Polyadenylated RNAs were then bound to Oligo-d(T)25 magnetic beads (NEB) in the presence of 0.5 M LiCl and treated with RppH (RNA pyrophosphohydrolase, NEB) to convert 5’ triphosphate moieties into single phosphate moieties. A 24 nt RNA adapter (CACACGCACACACAACCAGAGGAG) was then ligated to the 5’ ends of the product RNAs by T4 RNA ligase I (NEB) in the presence of 12.5% (v/v) PEG. RNAs were released from beads by heating to 75 °C for 2 min. RNAs were purified using Agencourt RNAClean XP beads, the sequencing library was prepared using the Direct RNA sequencing kit (Oxford nanopore) and sequenced using a MinION Mk1D instrument (Oxford nanopore).

The reference RNA sequences were constructed for each of the seven promoters assuming that transcription initiates at a purine separated by 6 or 7 bp from the −10 promoter element or at a purine separated by 7 bp from the −10 element if the former is not present. Reads (FASTQ format) were aligned to the reference RNAs using MinKNOW (BAM format output). The default alignment settings in MinKNOW are relaxed enough to recover reads with small indels at the adapter-RNA 5’ junction and the same reads were recovered if the reference sequences were constructed using transcription start sites ±2 nucleotides. Reads that overlap with the adapter sequence (first 24 nt) were extracted (BAM format output) and converted back to a FASTQ format list using SAMtools. Reads were then loaded into Ugene and searched with sequence patterns containing 6 nucleotides of the adapter and 7 nucleotides of RNA transcribed from the predicted transcription start site ±2 nucleotides. Additional patterns were designed after slippage was detected by manual inspection of read alignments. To perform the manual inspection, extracted reads were realigned to the reference RNAs using Muscle ^9^ to reveal sequence insertions that are obscured when viewing BAM format alignments.

#### *In vitro* crosslinking with DSSO followed by mass spectrometry analysis

*S. africana* holoenzyme (100 µl, 15 µM) with and without CarD (15 µM) was dialyzed against high-salt phosphate buffered saline (PBS; pH 7.4, 300 mM NaCl) resulting in 150 µl of 10 µM solution. Proteins were crosslinked with 3 mM disuccinimidyl sulfoxide (DSSO) for 30 min at 25 °C, quenched with Tris-HCl pH 7.9 (30 mM final) and dialyzed against PBS (pH 7.6, 150 mM NaCl). Samples were treated with urea, proteins were reduced with 10 mM dithiothreitol (in 50 mM Tris-HCl, pH 8.0), alkylated with 40 mM iodoacetamide and digested overnight with sequencing grade modified trypsin (Promega). Peptides were desalted using Sep-Pak tC18 96-well plate (Waters) and dried in a vacuum centrifuge. The dried peptide samples were dissolved with 0.1% formic acid and 1500 ng were subjected to the Liquid Chromatography-Electrospray Ionization-Tandem Mass Spectrometry (LC-ESI-MS/MS) analysis. The LC-ESI-MS/MS analyses were performed on a nanoflow HPLC system (Easy-nLC1000, Thermo Scientific) coupled to the Orbitrap Fusion Lumos mass spectrometer (Thermo Scientific, Bremen, Germany) equipped with a nano-electrospray ionization source and FAIMS Pro interface (Thermo Scientific). Compensation voltages of - 40 V, −60 V, and −75 V were used. Peptides were first loaded on a trapping column and subsequently separated inline on a 15 cm C18 column (75 μm x 15 cm, ReproSil-Pur 3 μm 120 Å C18-AQ, Dr. Maisch HPLC GmbH). The mobile phase consisted of water with 0.1% formic acid (solvent A) or acetonitrile/water (80:20 (v/v)) with 0.1% formic acid (solvent B). Peptides were eluted with the following gradient: from 5% to 21% of solvent B in 62 min, from 21% to 36% of solvent B in 48 min, from 36% to 100% of solvent B in 5 min, followed by 5 min at 100% of solvent B. MS data was acquired automatically by using Thermo Xcalibur 4.7 software (Thermo Fisher Scientific). Crosslinks were analyzed using the MS2-MS2-MS3 method where MS1 spectra were acquired in the Orbitrap mass analyzer with mass range of 375–1500 m/z at a resolution of 60,000. For the MS2-MS3-MS2 fragmentation method, sequential CID and ETD spectra were acquired for each precursor. MS3 scans were triggered by a targeted mass difference of 31.9721 detected in the MS2 scan. The MS3 scan was performed in the ion trap with CID fragmentation. Pairs of crosslinked peptides were identified by searching the raw data against the *S. africana* RNAP holoenzyme subunits using Proteome Discoverer 3.2 software (Thermo Scientific) and the XlinkX 3.2 algorithm (**Supplementary Data File**).

---

**Supplementary Table 1.**
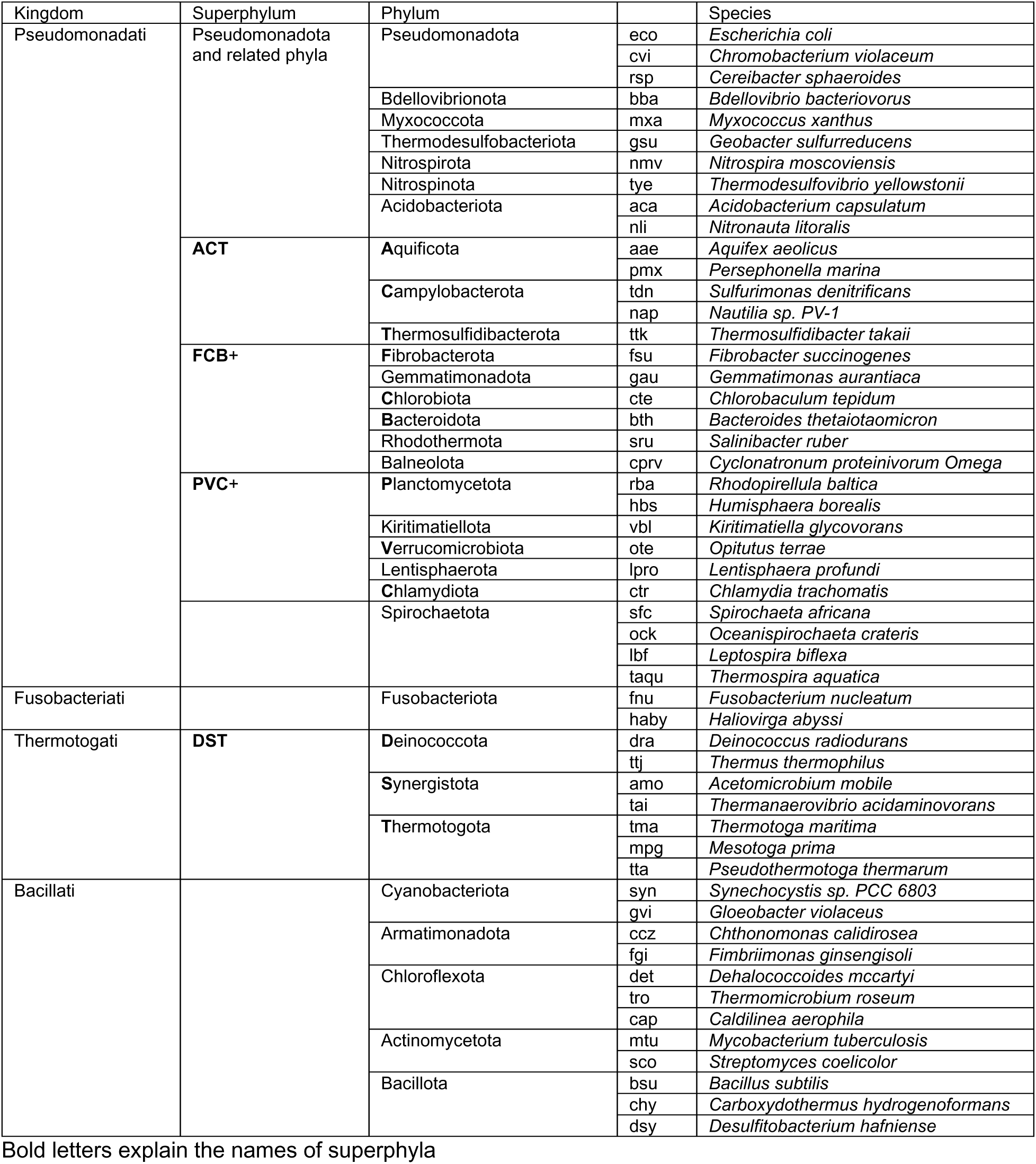
Bacterial species employed in analysis presented in Figure 1.

**Supplementary Table 2.**
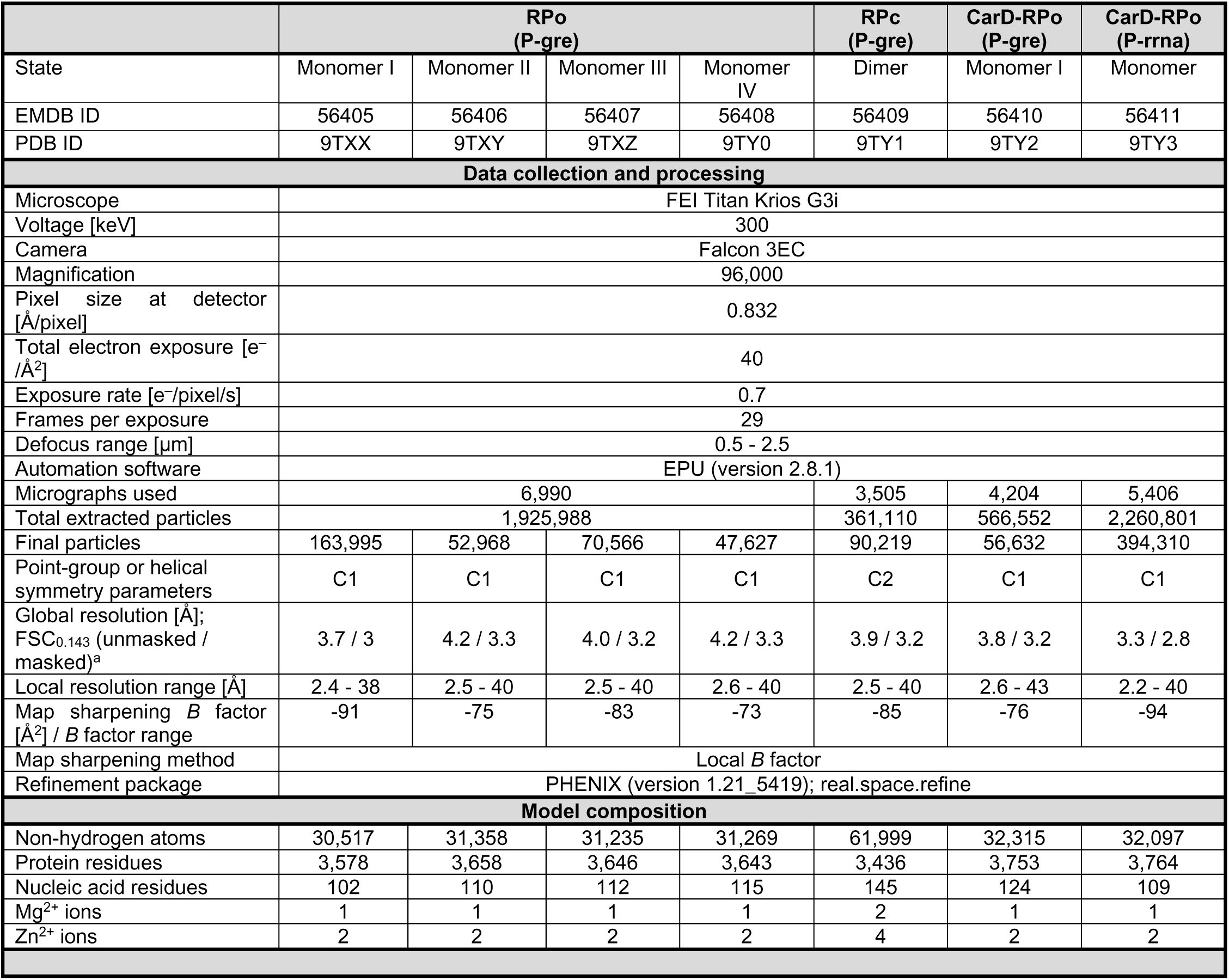
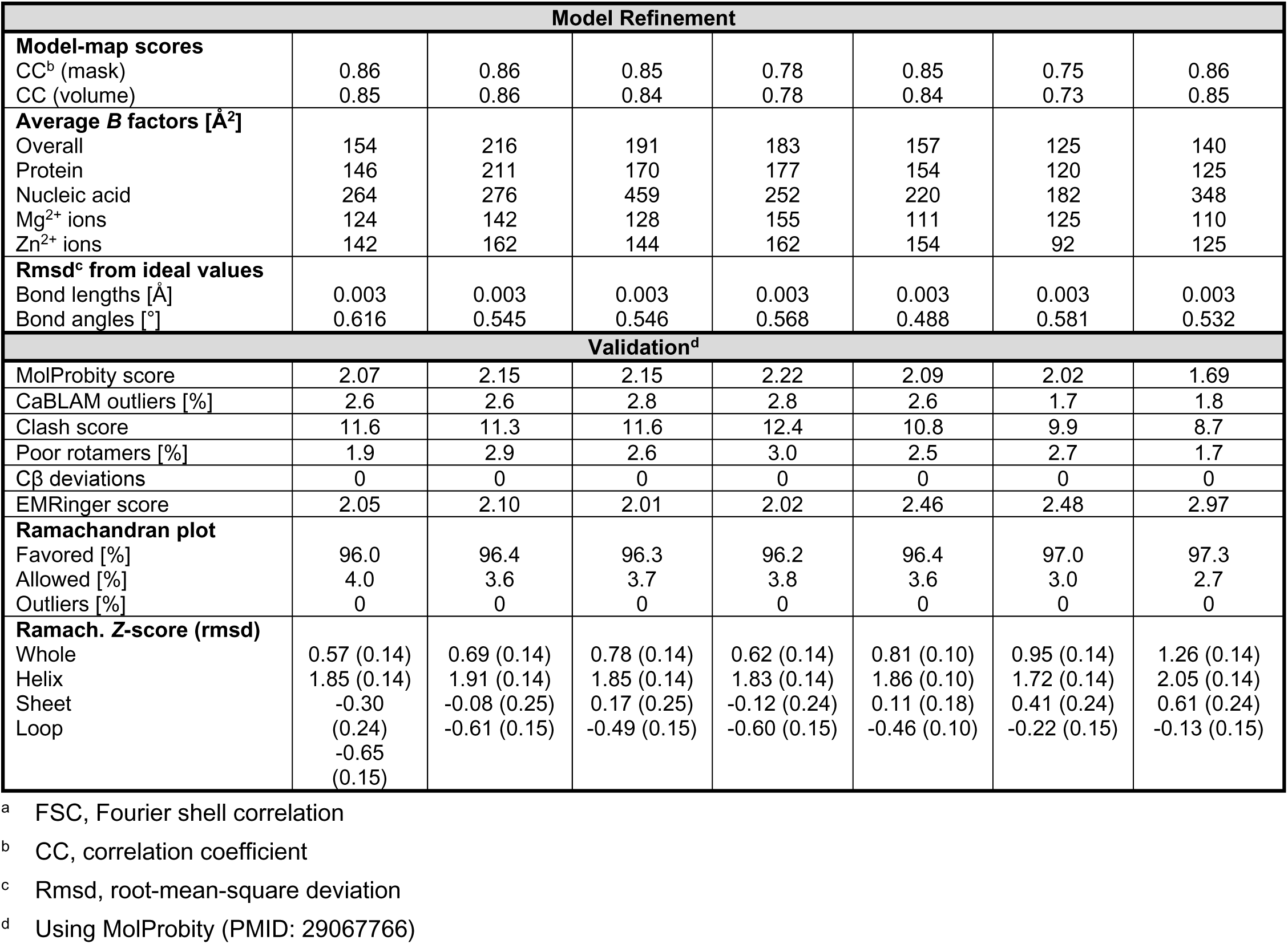
Cryo-EM data collection, refinement and validation statistics.

**Supplementary Table 3.**
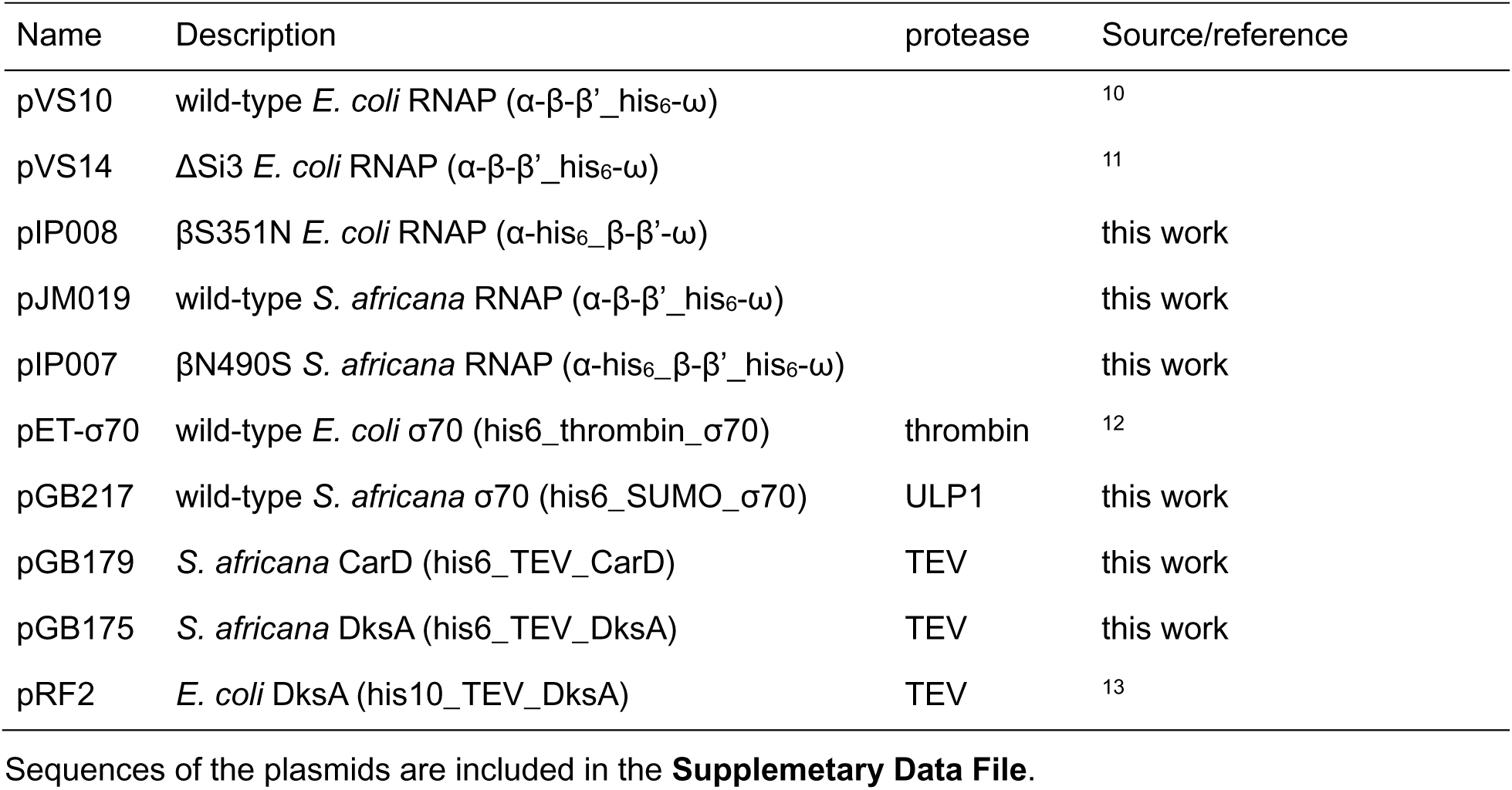
*E. coli* protein expression vectors used in this study.

**Supplementary Table 4.** Plasmids encoding transcription templates.

| Name | Description | downstream gene in the genome | Source/reference |
| --- | --- | --- | --- |
| pOP004 | P-gre with altered ITR | n/a | 14 |
| pVN003 | P-rrna promoter | spiaf_2142, spiaf_2147, spiaf_2596 | this work |
| pVN001 | P-gre promoter | spiaf_1527 | this work |
| pJK003 | P-prs promoter | spiaf_1522 | this work |
| pJK006 | P-trna promoter | spiaf_0540 | this work |
| pVL017 | P-fla promoter | spiaf_1672 | this work |
| pVL019 | P-pori promoter | spiaf_0679 | this work |
| pVL026 | promoter-less | n/a | this work |
| pJK004 | *E. coli* rrnB P1 | *E. coli* rRNA genes (16S, 23S, 5S) | this work |

Sequences of the plasmids are included in a **Supplementary Data File**.

**Supplementary Fig. 1.**
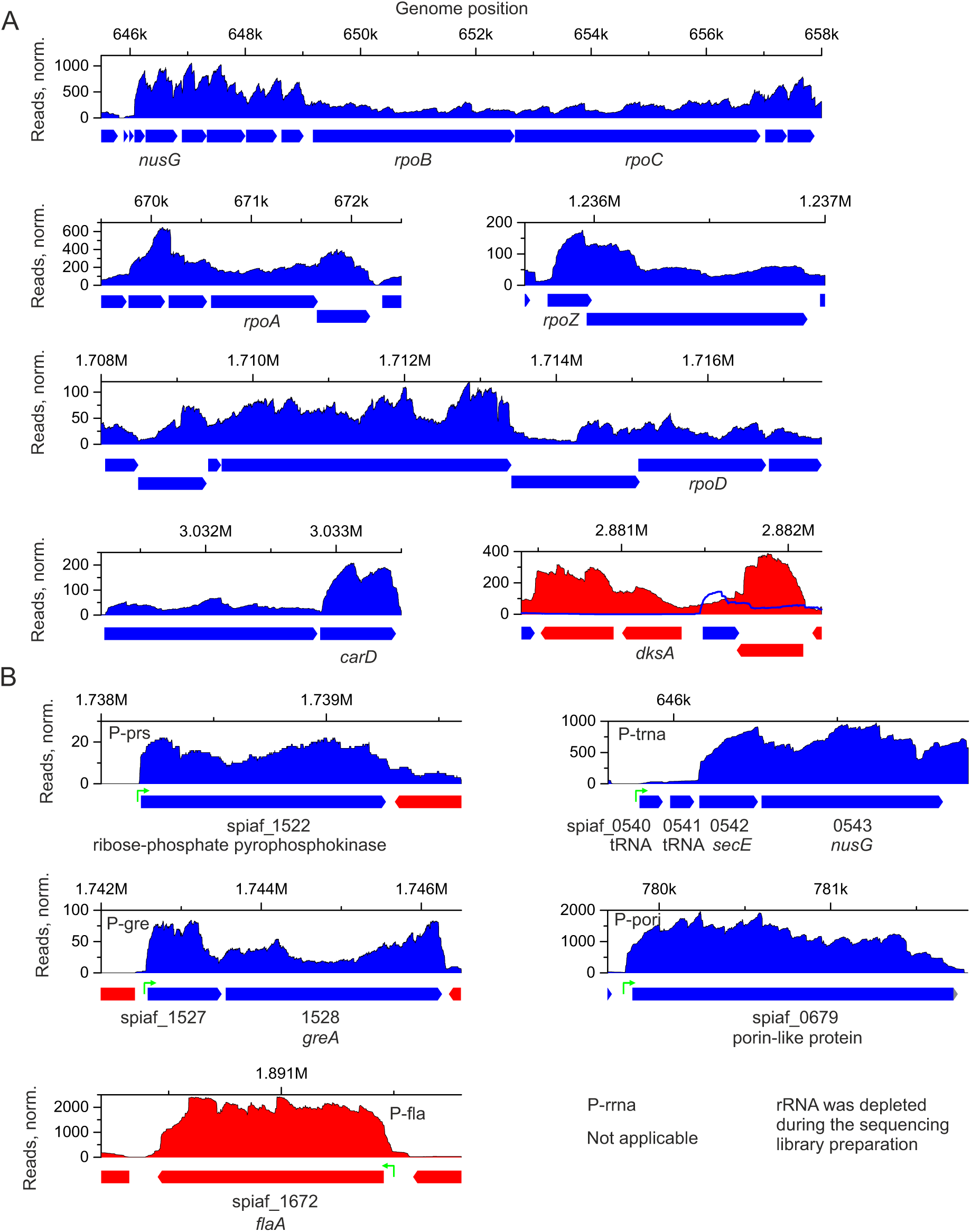
Strand-specific transcriptome coverage corresponding to the components of *S. africana* transcription system (A) and promoters investigated in this study (B). Annotated genes are plotted below the transcriptome graphs. KEGG (www.kegg.jp) gene numbers (spiaf_xxxx) and selected gene names are indicated below the annotations. Coloring: positive strand coverage and annotations blue, negative strand coverage and annotations red.

**Supplementary Fig. 2.**
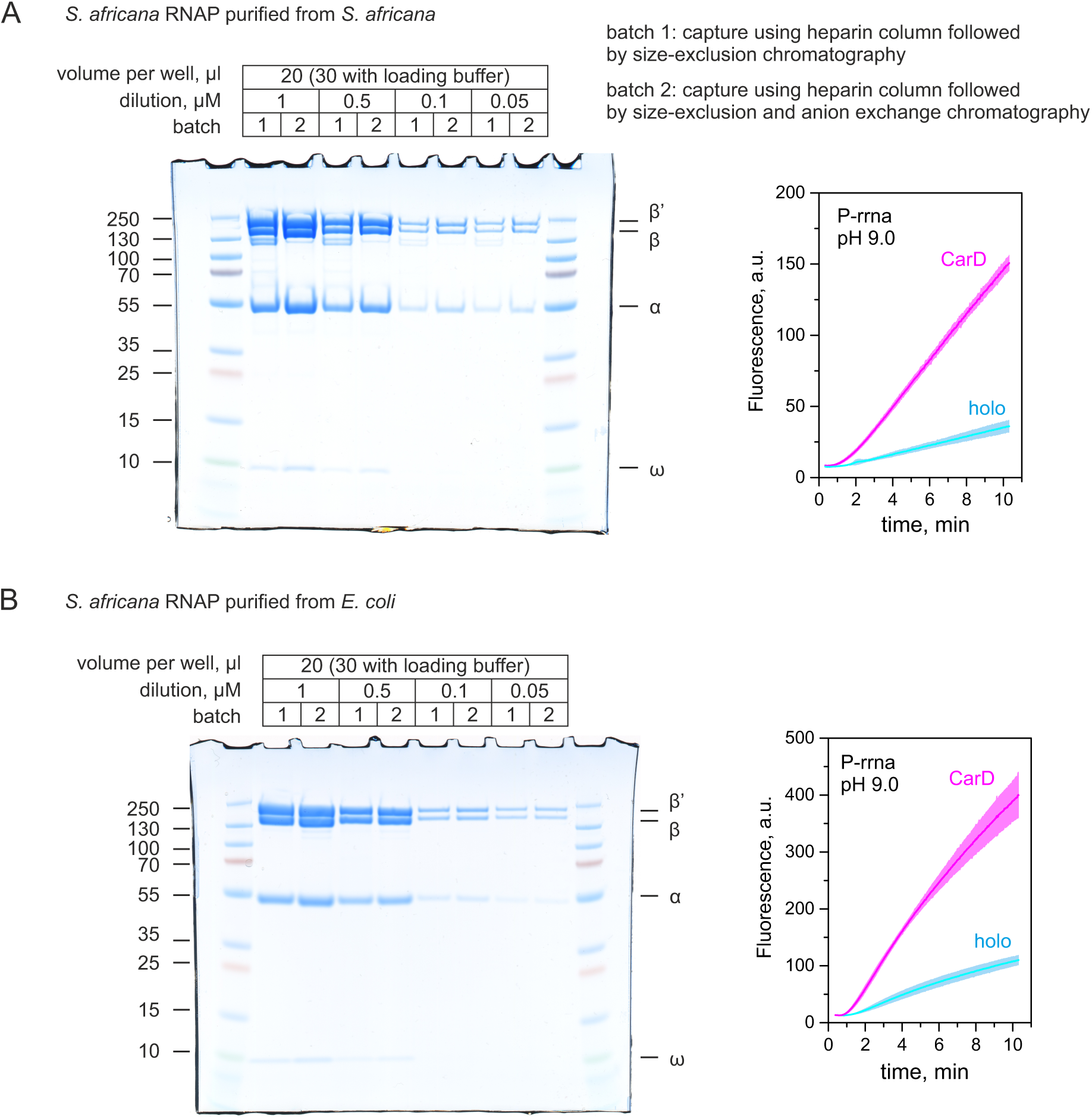
Transcriptional activity of *S. africana* RNAP purified from *S. africana* (A) and *E. coli* (B) at P-rrna promoter. SDS-PAGE analysis was performed using the MES buffer system (50 mM MES, 50 mM Tris Base, 0.1% SDS, 1 mM EDTA, pH 7.3). Transcription activity was monitored using FLAP assay (1 µM RNAP, 4 µM σ, 50 nM DNA template and where indicated 10 µM CarD). Coloring: holoenzyme (cyan), in the presence of CarD (magenta). Promoter sequences and schematic of the FLAP assay are presented in the main text Figure 2.

**Supplementary Fig. 3.**
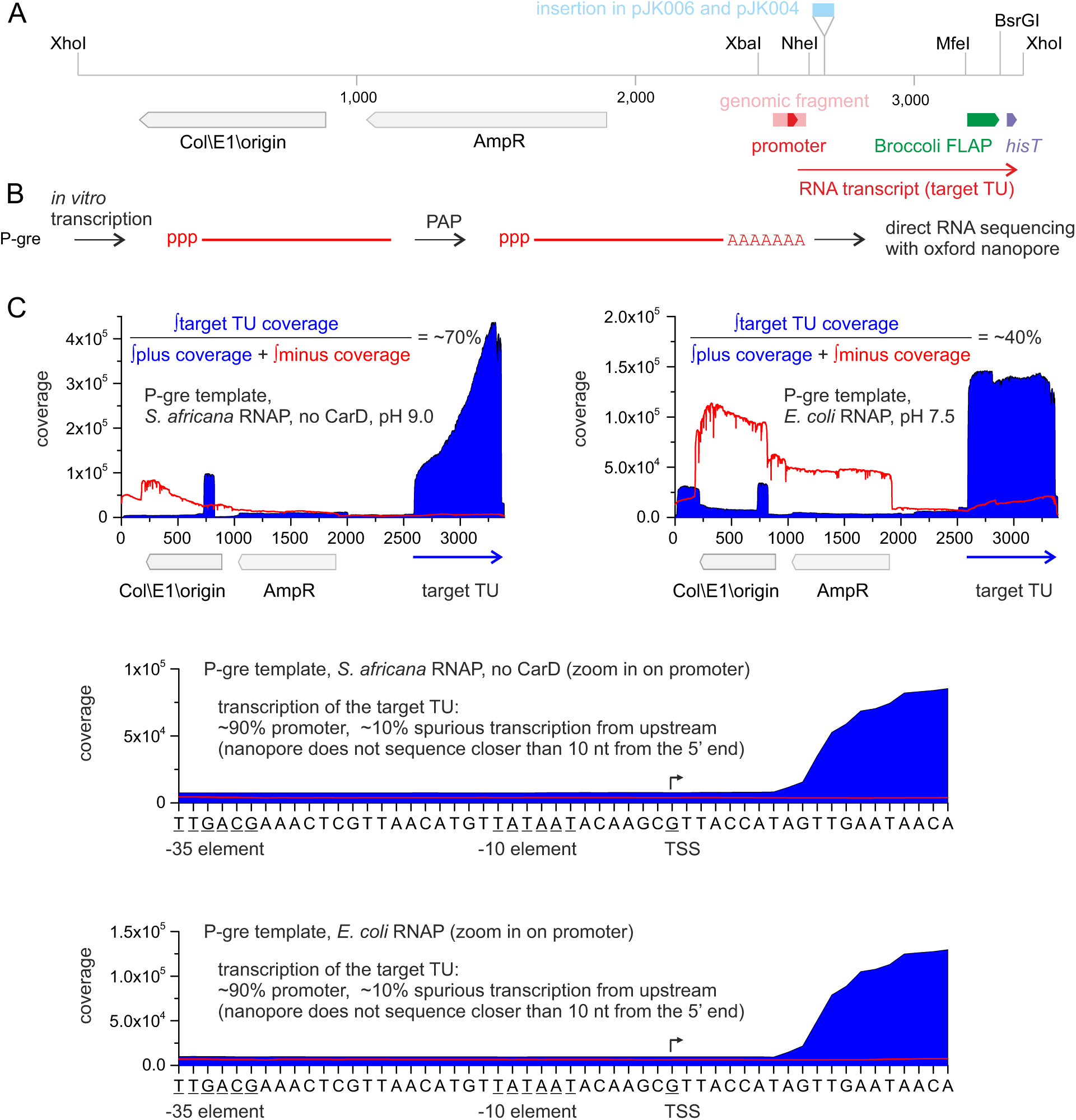
Analysis of RNA products transcribed by *S. africana* and *E. coli* holoenzymes from a linearized plasmid using direct RNA sequencing. **(A)** Schematics of transcription templates used in this study. pVN001, pVN003, pJK003, pVL017, pVL019 and pOP005 plasmids differed only in the *Xba*I-*Nhe*I module containing plasmid-specific fragment of *S. africana* genome (pink) including the promoter (red). pJK006 additionally contained an insertion in the *Nhe*I-*Mfe*I module. *Mfe*I-*BsrG*I module encoded Broccoli FLAP embedded in tRNA scaffold, *BsrG*I-*Xho*I module encoded intrinsic transcription terminator. **(B)** Schematics of the experiment. *In vitro* transcription reactions (0.5 ml) were performed as in FLAP assays. Product RNA were purified using RNA easy kit (Qiagen), polyadenylated with *E. coli* poly-A polymerase (PAP). The sequencing library was prepared using Direct RNA sequencing kit (Oxford nanopore), and sequenced using MinION Mk1D instrument. **(C)** Reads were aligned to the P-gre transcription template using MinKNOW software, alignment files were split according to the strand using SAMtools, loaded into Ugene, per base coverage was exported and plotted using Origin software.

**Supplementary Fig. 4.**
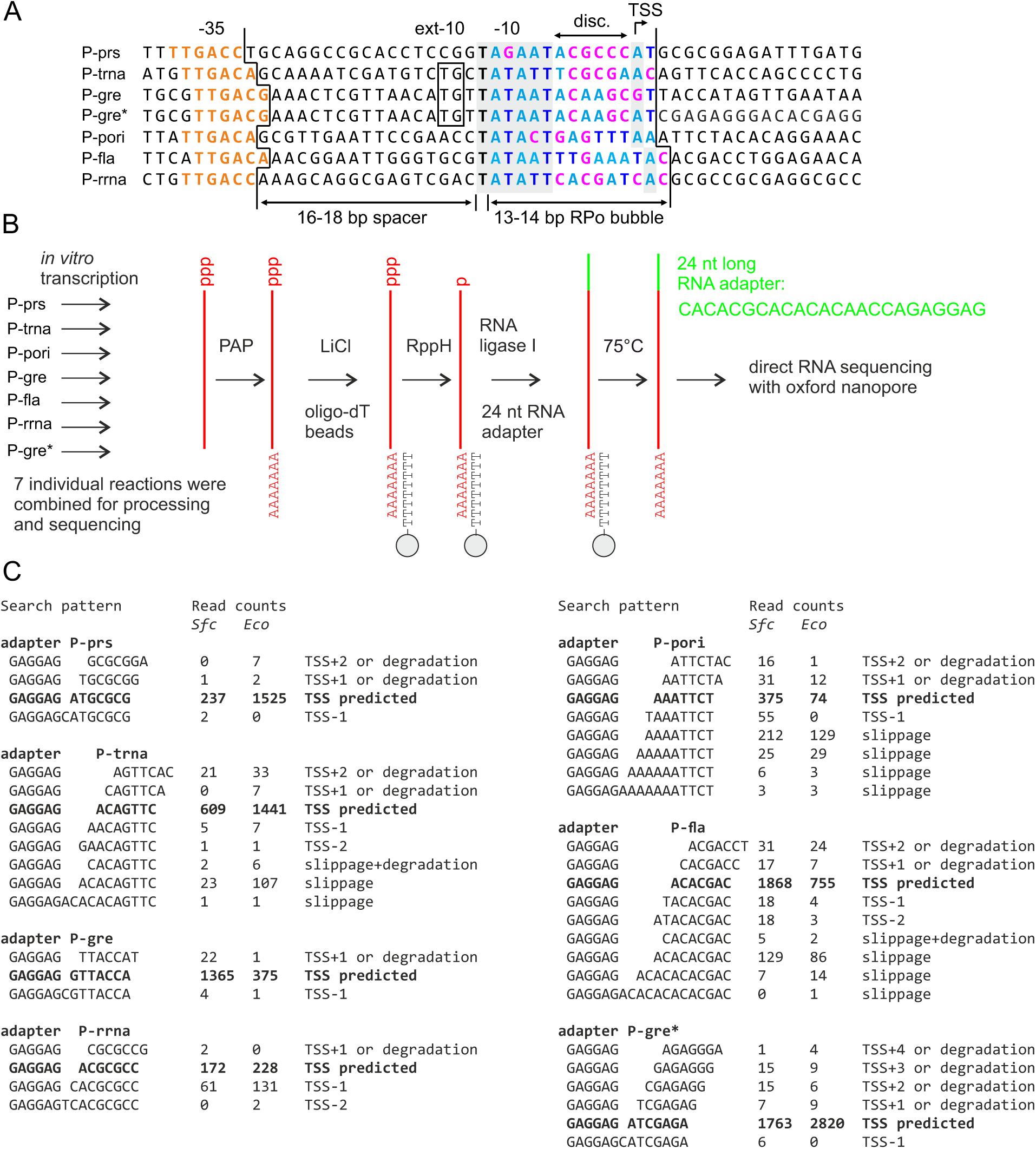
Transcription start site mapping using 5’ adapter ligation followed by direct RNA sequencing. **(A)** Promoter sequences used in this study aligned by −10 elements. Color coding as in main text Fig. 2. **(B)** The schematics of *in vitro* RNA processing before sequencing. **(C)** Counts of reads that map to the adapter-RNA 5’ end junction and contain indicated sequence patterns. RNAs were transcribed individually from 7 templates and combined for processing and sequencing. Separate sequencing runs were performed for *S. africana* and *E. coli* holoenzymes. Search patterns contain 6 nucleotides of the adapter and 7 nucleotides of RNA transcribed from the predicted TSS ±2 nucleotides. Additional patterns were designed after slippage was detected by manual inspection of read alignments. Patterns and read counts corresponding to the predicted TSS are indicated in bold text. RNA sequencing produced U-containing reads, but MinKNOW aligner converted U to T. Sequence patterns were then designed to feature T in place of U.

**Supplementary Fig. 5.**
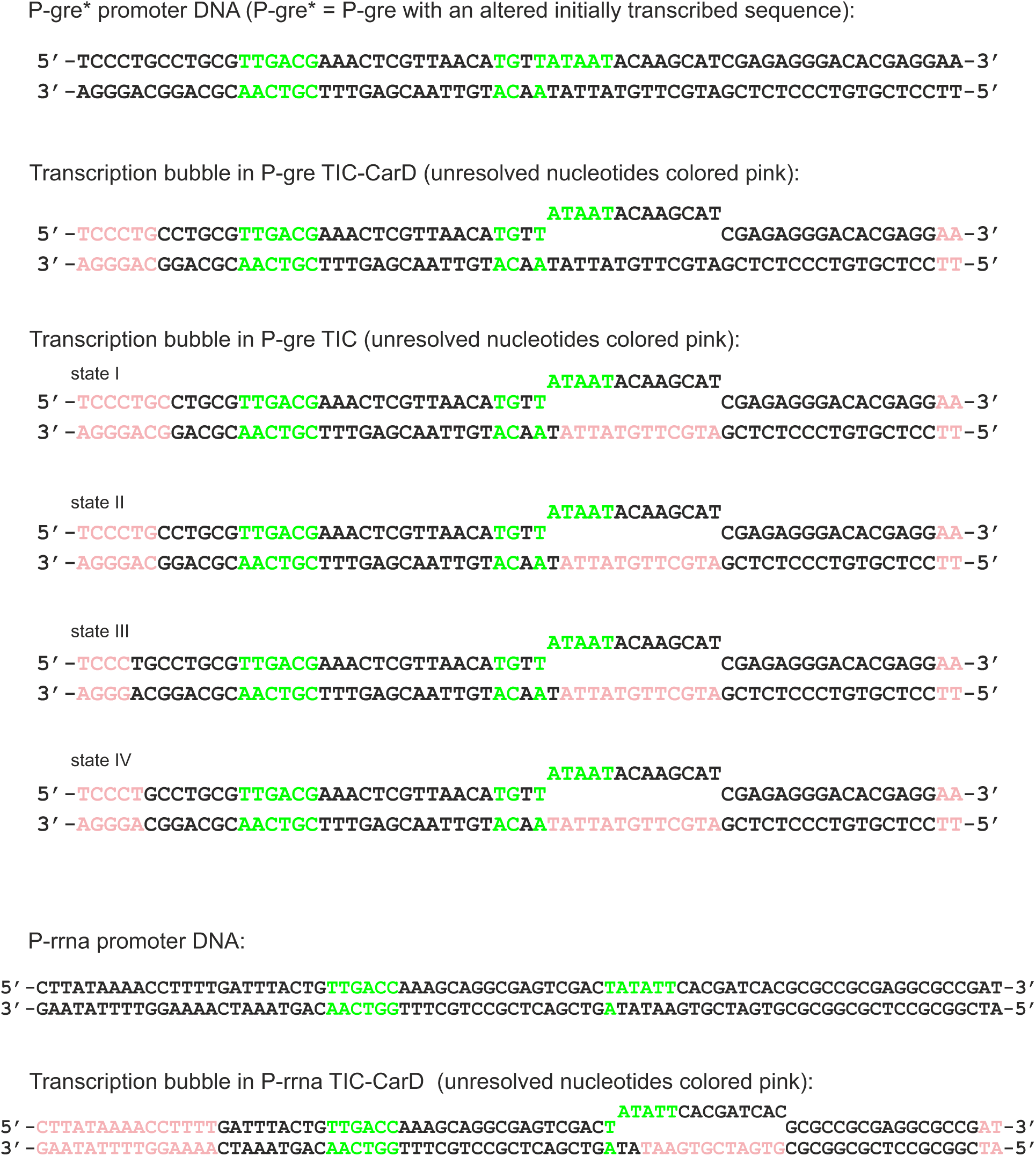
Promoter scaffolds used to assemble *S. africana* holoenzyme - DNA complexes for cryoEM analysis. Promoter elements are shaded in green. Nucleotides that were not represented by the density and omitted from structural models are shaded light pink.

**Supplementary Fig. 6.**
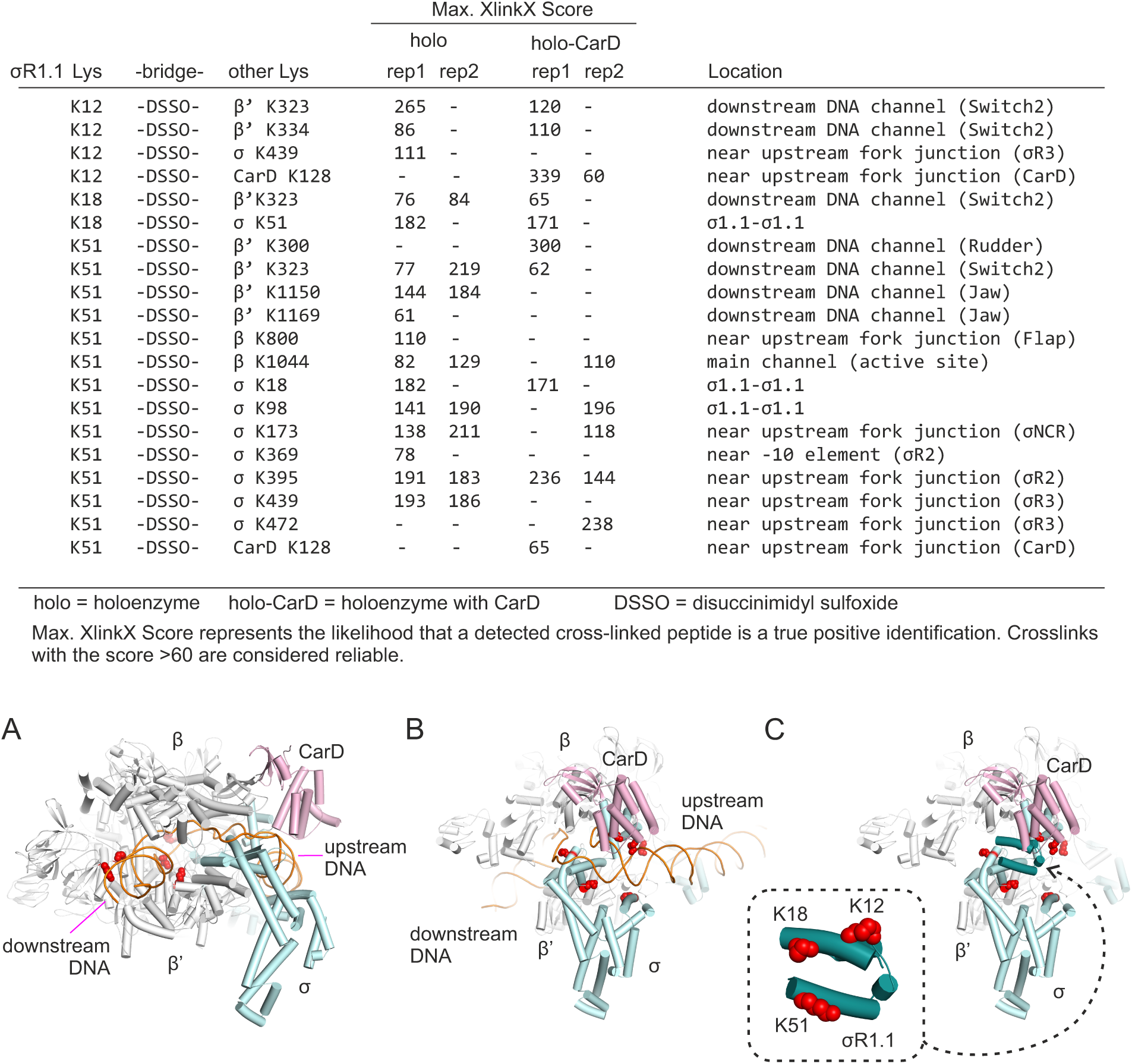
Probing the location of σR1.1 in *S. africana* holoenzyme by *in vitro* crosslinking with disuccinimidyl sulfoxide (DSSO) followed by mass spectrometry analysis. Crosslinks involving Lys residues within the globular domain of σR1.1 (top table) were manually selected from the pool of all identified crosslinks (**Supplementary Data File**). Lys residues involved in crosslinks with σR1.1 lysines were then visualized (red spheres) on the structure of the *S. africana* RPo at the P-gre promoter with CarD. **(A)** σR1.1-crosslinkable Lys residues in Switch 2 region, the Rudder Loop and the Jaw domain suggest that σR1.1 is located in the downstream DNA channel. **(B)** σR1.1-crosslinkable Lys residues in σR2, σR3, σNCR and CarD suggest that σR1.1 is located in place of the upstream fork junction. **(C)** Same as in (B) but the promoter DNA was removed from the RPo structure and σR1.1 from the dimeric holoenzyme structure was modelled in using β subunit as a reference for structural alignment. The inset shows the zoomed in σR1.1 and σR1.1 Lys residues (red spheres) that form crosslinks discussed above.

**Supplementary Fig. 7.**
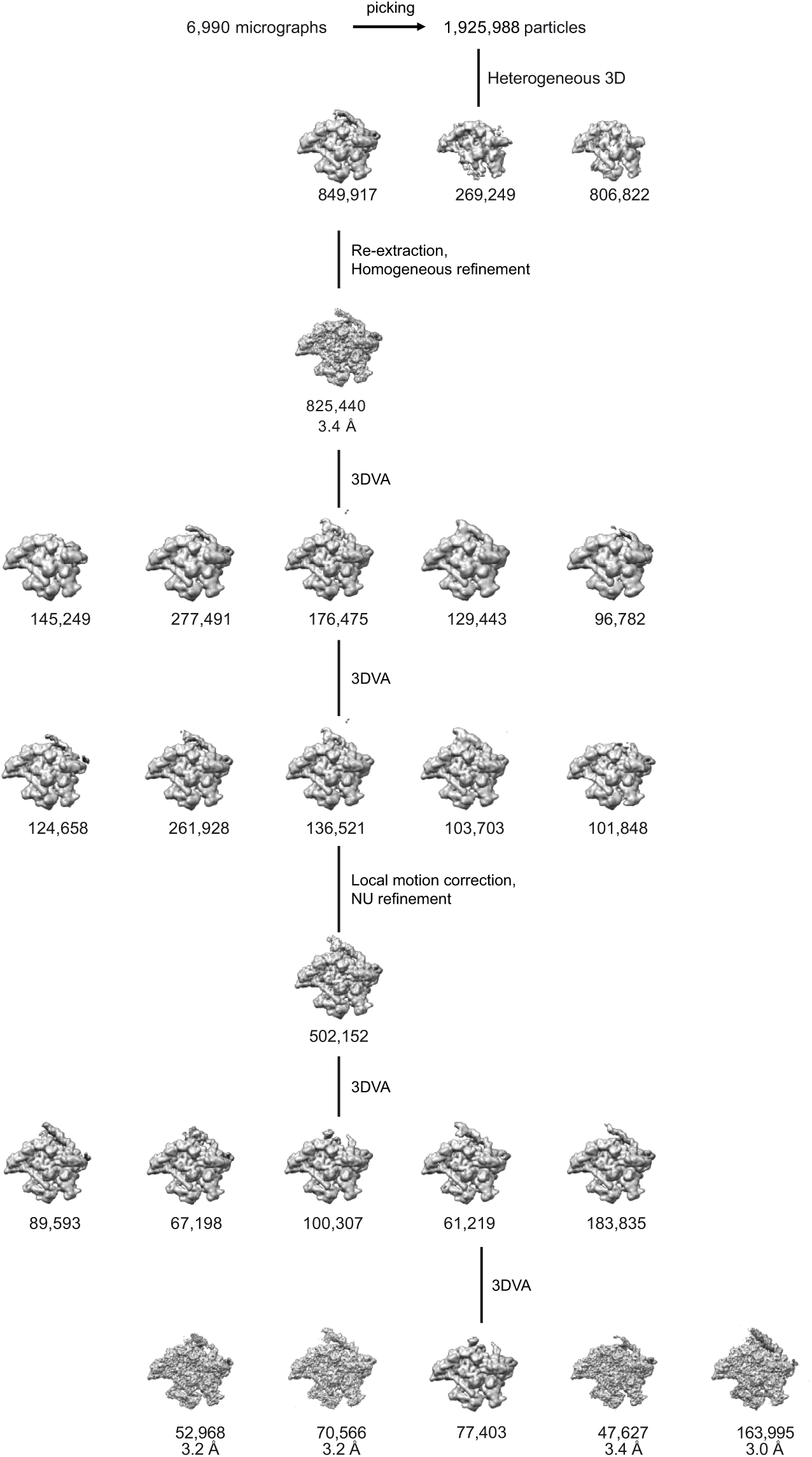
**Hierarchical clustering analysis of particle images of monomeric *S. africana* holoenzyme-DNA complex assembled at P-gre promoter**

**Supplementary Fig. 8.**
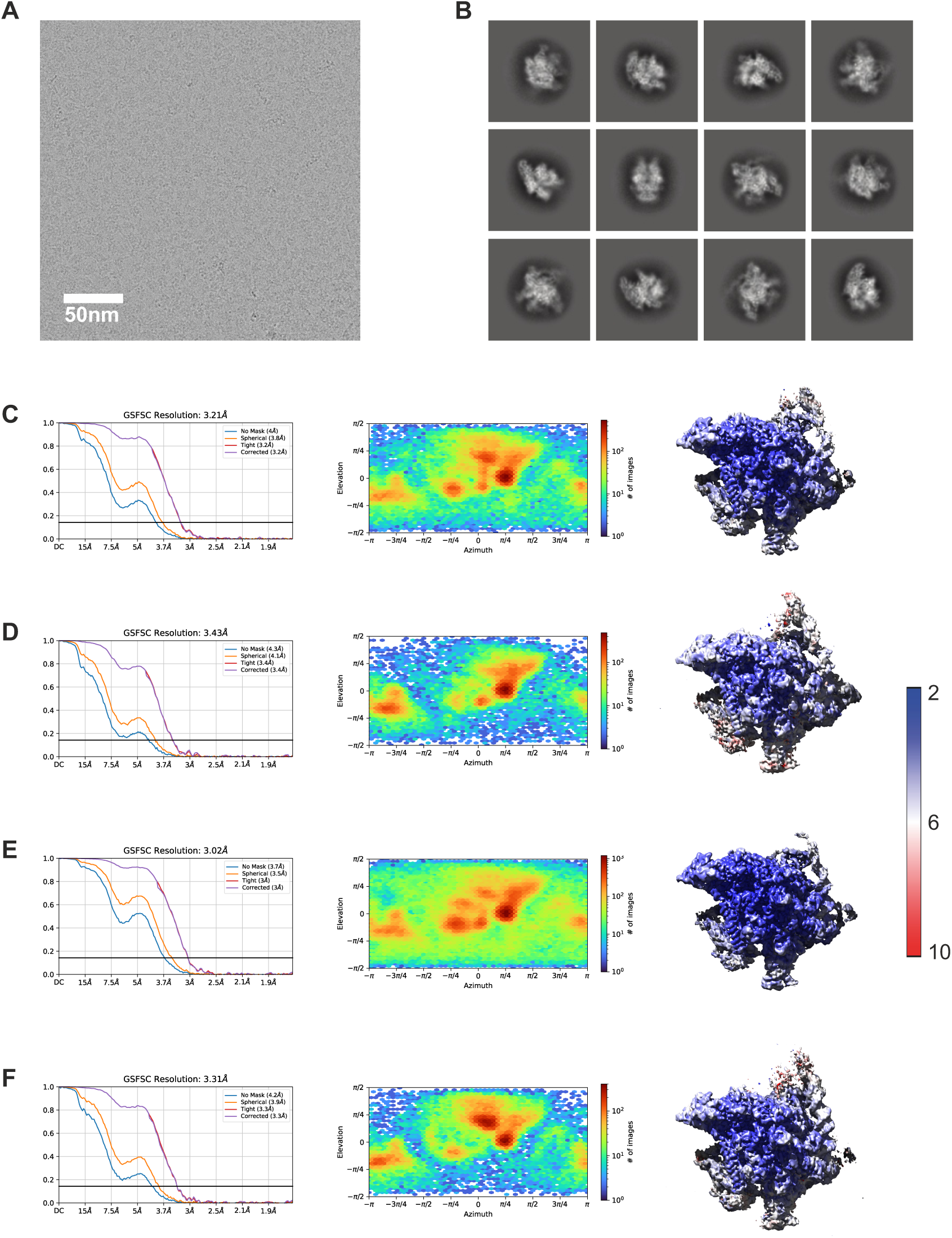
CryoEM data processing, monomeric *S. africana* holoenzyme-DNA complex assembled at P-gre promoter. **(A)** Representative cryo-electron micrograph of the particles. Scale bar, 50 nm. **(B)** 2D class averages of the particle images. **(C-F)** Left, Fourier shell correlation (FSC) plots for half-maps of four states of the monomeric complex with 0.143 FSC criteria indicated; nominal resolutions 3.2 Å, 3.4 Å, 3.0 Å and 3.3 Å. Middle, the distributions of particle orientations show preferential orientation bias. Right, local resolution plots.

**Supplementary Fig. 9.**
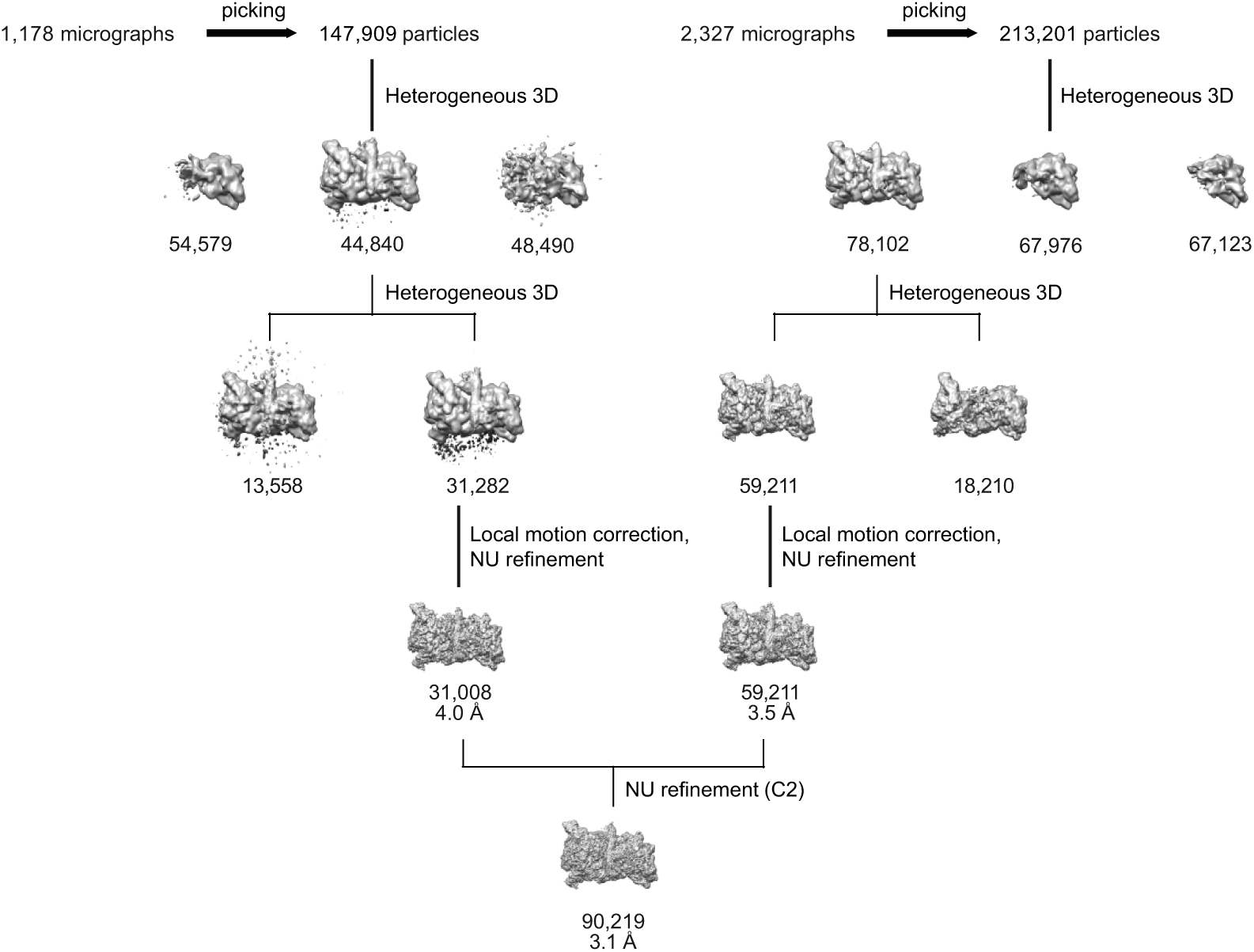
**Hierarchical clustering analysis of particle images of dimeric *S. africana* holoenzyme-DNA complex assembled at P-gre promoter.**

**Supplementary Fig. 10.**
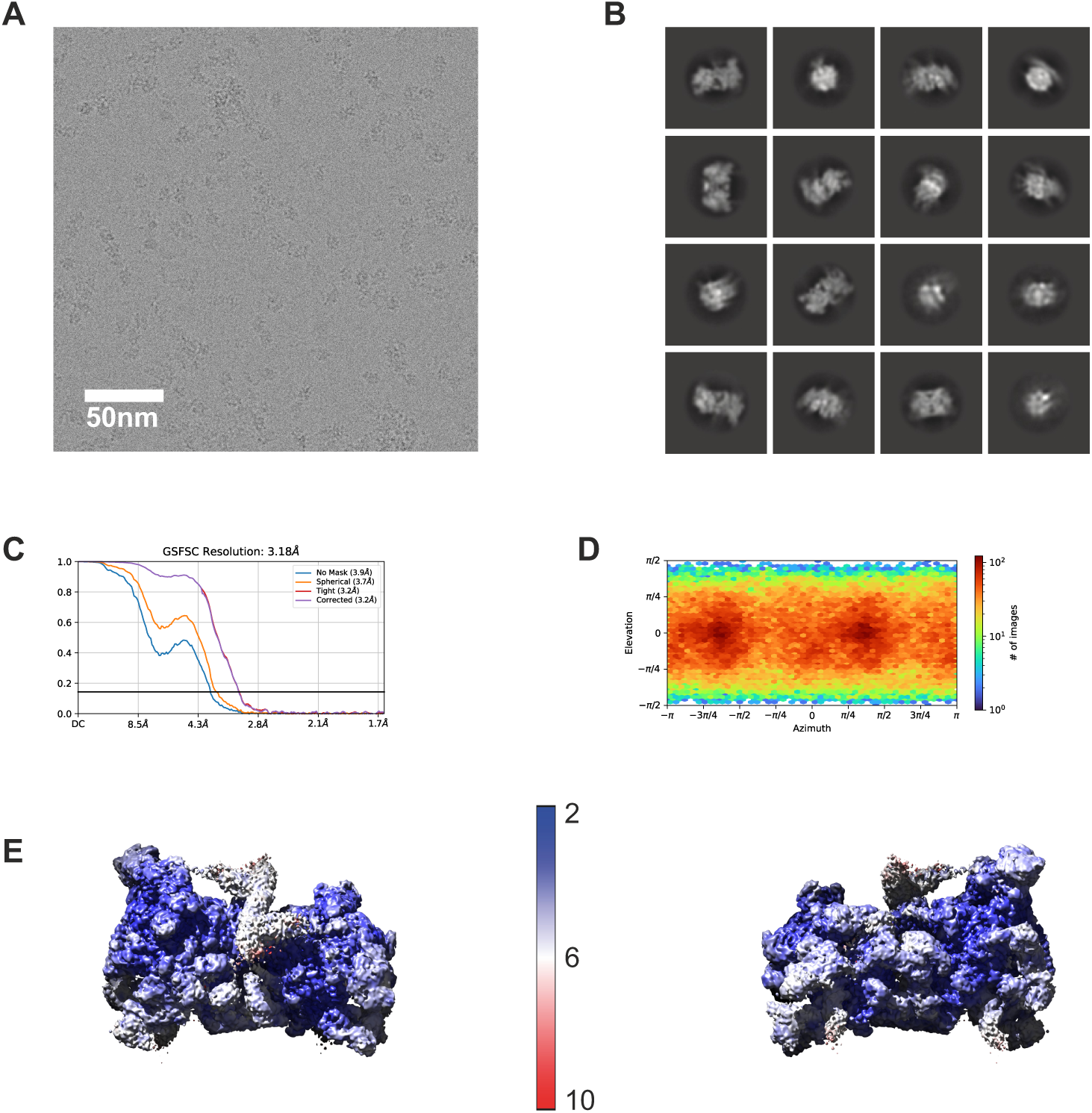
CryoEM data processing, dimeric *S. africana* holoenzyme-DNA complex assembled at P-gre promoter DNA. **(A)** Representative cryo-electron micrograph of the particles. Scale bar, 50 nm. **(B)** 2D class averages of the particle images. **(C)** Fourier shell correlation (FSC) plot for half-maps of the dimeric complex with 0.143 FSC criteria indicated; nominal resolution 3.2 Å. **(D)** The distribution of particle orientations shows preferential orientation bias. **(E)** Local resolution plots.

**Supplementary Fig. 11.**
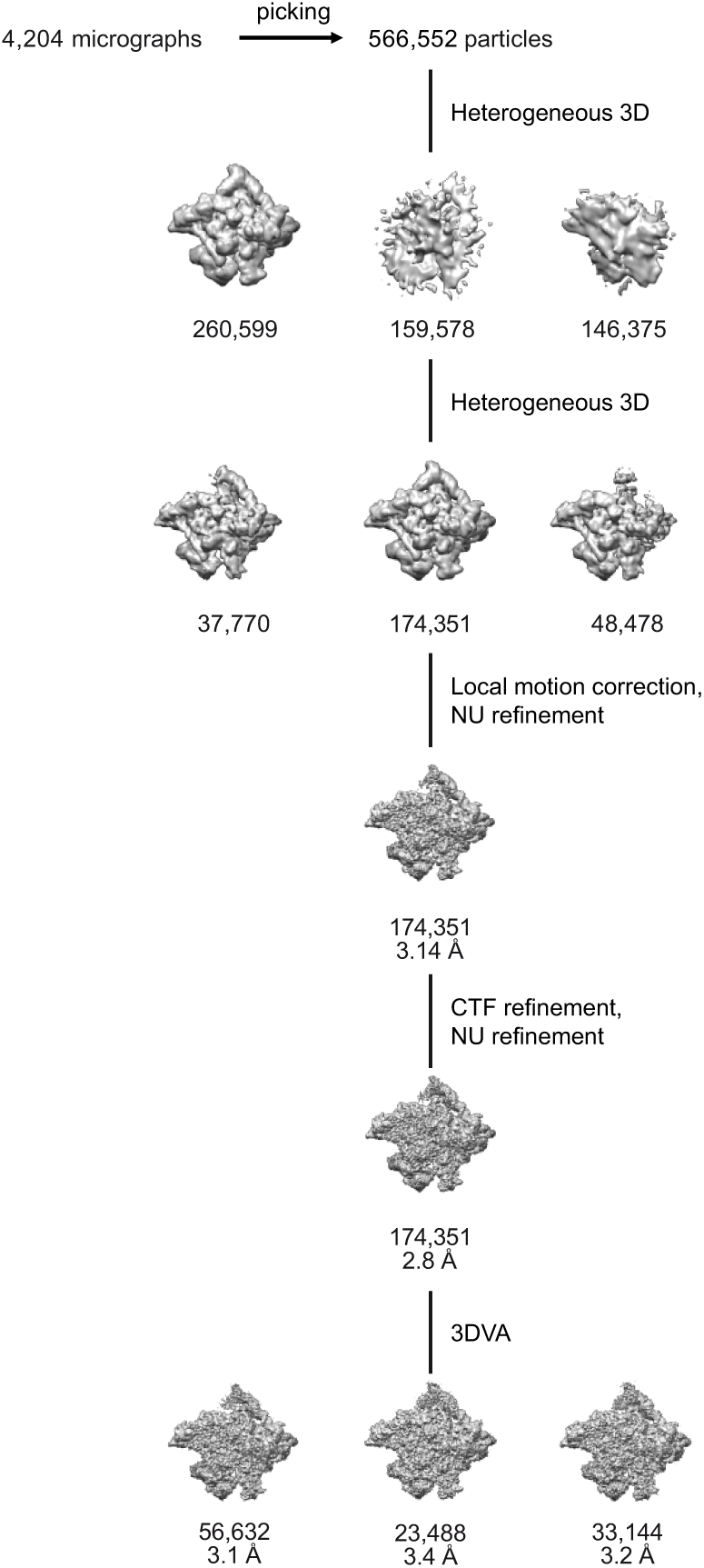
**Hierarchical clustering analysis of particle images of CarD-modified *S. africana* holoenzyme-DNA complex assembled at P-gre promoter.**

**Supplementary Fig. 12.**
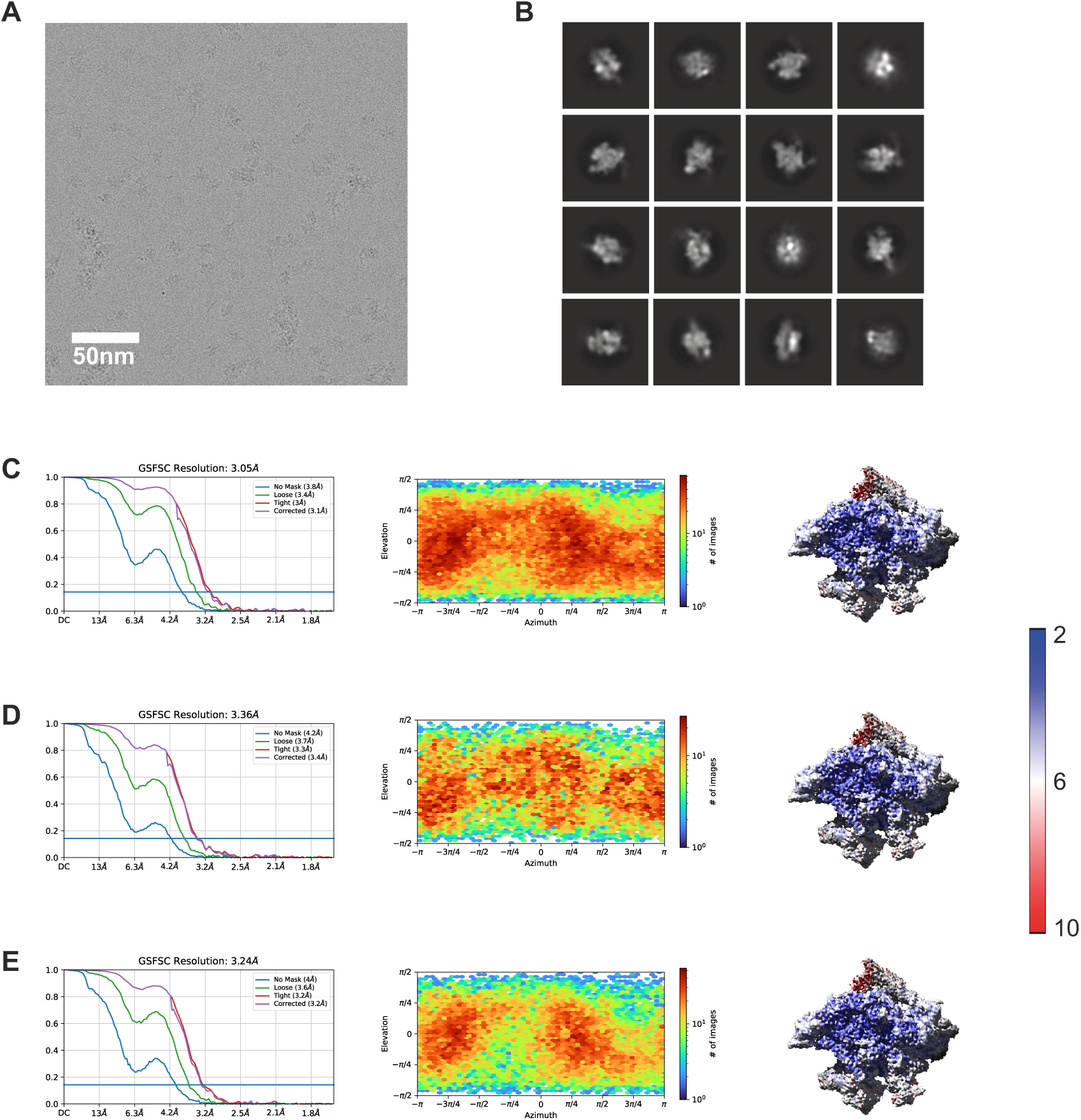
CryoEM data processing, CarD-modified *S. africana* holoenzyme-DNA complex assembled at P-gre promoter. **(A)** Representative cryo-electron micrograph of the particles. Scale bar, 50 nm. **(B)** 2D class averages of the particle images. **(C-E)** Left, Fourier shell correlation (FSC) plots for half-maps of three states of the complex with 0.143 FSC criteria indicated; nominal resolutions 3.0 Å, 3.3 Å and 3.2 Å. Middle, the distributions of particle orientations show preferential orientation bias. Right, local resolution plots.

## Notes

### Competing Interest Statement

The authors have declared no competing interest.

### Summary of Updates

ORCID IDs added for all authors jPOST link and accession code added Minor formatting changes

